# Sugar co-ordinates plant defence signalling

**DOI:** 10.1101/2023.07.20.549975

**Authors:** Kohji Yamada, Akira Mine

**Affiliations:** Graduate School of Technology, Industrial and Social Sciences, Tokushima University, Tokushima, Japan; JST, PRESTO, Kawaguchi, Japan; Graduate School of Agriculture, Kyoto University, Kyoto, Japan

## Abstract

Recognition of microbial molecules triggers energy-intensive defence systems. Although successful defence should therefore depend on energy availability, whether and, if so, how cellular metabolic information is molecularly input into defence remains unclear. We show that sugar, especially glucose-6-phosphate (G6P), plays a key role in regulating the types and amplitudes of defence outputs in *Arabidopsis thaliana*. Under sugar-sufficient conditions, protein and phosphorylation levels of calcium-dependent protein kinase 5 (CPK5) are elevated by induced expression and G6P-mediated suppression of protein phosphatases, priming defence responses. Furthermore, recognition of bacterial flagellin activates sugar transporters, leading to increased cellular G6P, which elicits CPK5-independent signalling promoting synthesis of the phytohormone salicylic acid (SA) involved in anti-bacterial defence. In contrast, while perception of fungal chitin does not promote sugar influx or SA accumulation, chitin-induced synthesis of the anti-fungal compound camalexin requires basal sugar influx activity. These results suggest that, by monitoring G6P levels, plants determine defence priming levels and execute appropriate outputs against bacterial and fungal pathogens. Together, our findings provide a comprehensive view of the roles of sugar in plant defence.

## Introduction

In plants, perception of pathogen-associated molecular patterns (PAMPs) on the plasma membrane triggers a cascade of defence responses, including extensive transcriptional reprogramming and *de novo* synthesis of metabolites such as phytoalexins and phytohormones (Couto and Zipfel, 2016). Despite the energy-intensive nature of defence signalling, the molecular link between defence signalling and cellular metabolic status remains largely unknown. For example, defence signalling should be suppressed under low nutritional conditions and enhanced under sufficient nutritional conditions. Of primary metabolites, sugar is known to affect various signal pathways not only as an energy precursor and a carbon skeleton but also as a signal molecule (Jang and Sheen, 1994). Sugar has been proposed to stimulate plant defence, by a mechanism whose molecular basis has remained unclear for over two decades (Sheen et al., 1999). In addition, exogenous application of sugars to plants is reported to enhance defence responses against bacterial and fungal pathogens (Gómez-Ariza et al., 2007; Tsutsui et al., 2015). Glucose (Glc) is recognized by hexokinase 1(HXK1) (Moore et al., 2003), a dual-function enzyme with both catalytic and sensory activity for Glc, in the model plant *Arabidopsis thaliana*. Sensing of Glc via HXK1 triggers down-regulation of photosynthetic genes (Moore et al., 2003). On the other hand, Glc catalysis mediated by HXK1 induces defence signalling. Overexpression of not only *Arabidopsis* HXK1 but also yeast HXK2 elevated expression of defence-responsive genes (Xiao et al., 2000). Furthermore, cell death induced from loss of the *myo*-inositol 1-phosphatase synthase 1 gene was attenuated in the absence of HXK1’s enzymatic activity (Bruggeman et al., 2015). Therefore, a Glc-derived metabolite(s) catalysed by HXK, rather than Glc *per se*, is thought to stimulate defence signalling (Sheen et al., 1999).

In this study, we focused on how cellular sugar conditions are molecularly input into plant defence signalling. We showed that Glc-6-phosphate (G6P), not Glc, affects multiple signal branches that orchestrate defence outputs, using genetic and pharmacological analyses. First, calcium-dependent protein kinase 5 (CPK5), which stimulates expression of sugar-responsive genes, is suppressed by clade A protein phosphatase 2Cs (PP2Cs). We found that G6P inhibits their phosphatase activities, leading to de-suppression of CPK5 activity. Therefore, defence responses are primed under sugar-sufficient conditions. Furthermore, after perception of bacterial and fungal PAMPs, sugar influx transporters regulate cellular G6P amounts, which elicits CPK5-indepdent signal pathways that co-operatively or antagonistically act on CPK5-mediated signalling for executing appropriate defence outputs against bacterial and fungal pathogens. Thus, this work provides a comprehensive view of the roles of sugar in co-ordinating plant defence signalling.

## Results

### Sugar activates expression of defence-related genes via HXK-mediated Glc phosphorylation

We first demonstrated that the application of sucrose (Suc) to *Arabidopsis* seedlings induced defence-related genes by RNA-seq analysis (Figure S1A, S1B, cluster III). In addition, we showed that Glc application also induced expression of the defence marker gene *NDR1/HIN1-LIKE 10* (*NHL10*) and the camalexin synthetic gene *PHYTOALEXIN DEFICIENT 3* (*PAD3*) (Figure S1C, S1D), showing that sugar stimulates defence signalling in *Arabidopsis*. On the other hand, expression of the photosynthesis-related gene *CHLOROPHYLL A/B BINDING PROTEIN 1* (*CAB1*) was repressed (Figure S1C, S1D), which confirmed the activation of sugar signalling under these experimental conditions (above 25 mM of Glc or Suc). We next investigated whether sugar also promotes gene expression during pattern-triggered immunity (PTI). Seedlings were treated with flg22 peptide, a derivative of bacterial flagellin, for 8 h with or without Glc, Suc or mannitol (Mtl) (Figure 1A). Flg22-triggered expression of *NHL10* and *PAD3* was higher in the presence of Glc or Suc, but unaffected by Mtl (Figure 1B). These data suggested that the recognition of metabolizable sugars and/or sugar metabolism affects defence signalling, and this effect is not due to alteration of osmotic pressure. Since Suc effects were sometimes stronger than Glc effects as for *PAD3* expression (Figure 1B), we next asked whether Glc and Suc differentially activate defence signalling. In *Arabidopsis*, two cytoplasmic HXKs, HXK1 and HXK2 (Moore et al., 2003), phosphorylate Glc but not Suc. To explore the involvement of HXKs in Glc- and/or Suc-mediated stimulation of defence signalling, we established *hxk1 hxk2* (*hxk1/2*) mutants (Figure S2F). The *hxk2* mutant was presumed to express HXK2 with a C-terminal 13-aa deletion (Figure S2C). The activity of HXK2ΔC13 was significantly weaker than that of HXK2 *in vitro* (Figure S2D). Therefore, the *hxk2* mutant was used in this study although HXK2 activity was not completely absent. We found that both Glc- and Suc-induced expression of *NHL10* and *PAD3* were reduced in *hxk1/2* plants (Figure 1C), suggesting that Glc, hydrolysed from Suc, stimulates their expression under Suc conditions. Suc accumulates to high levels in the cytoplasm of plant cells, while Glc is mainly compartmentalized in the vacuole (Leidreiter et al., 1995). Therefore, Suc may stimulate gene expression more easily than Glc, although both sugars act through HXKs.

**Figure 1.**
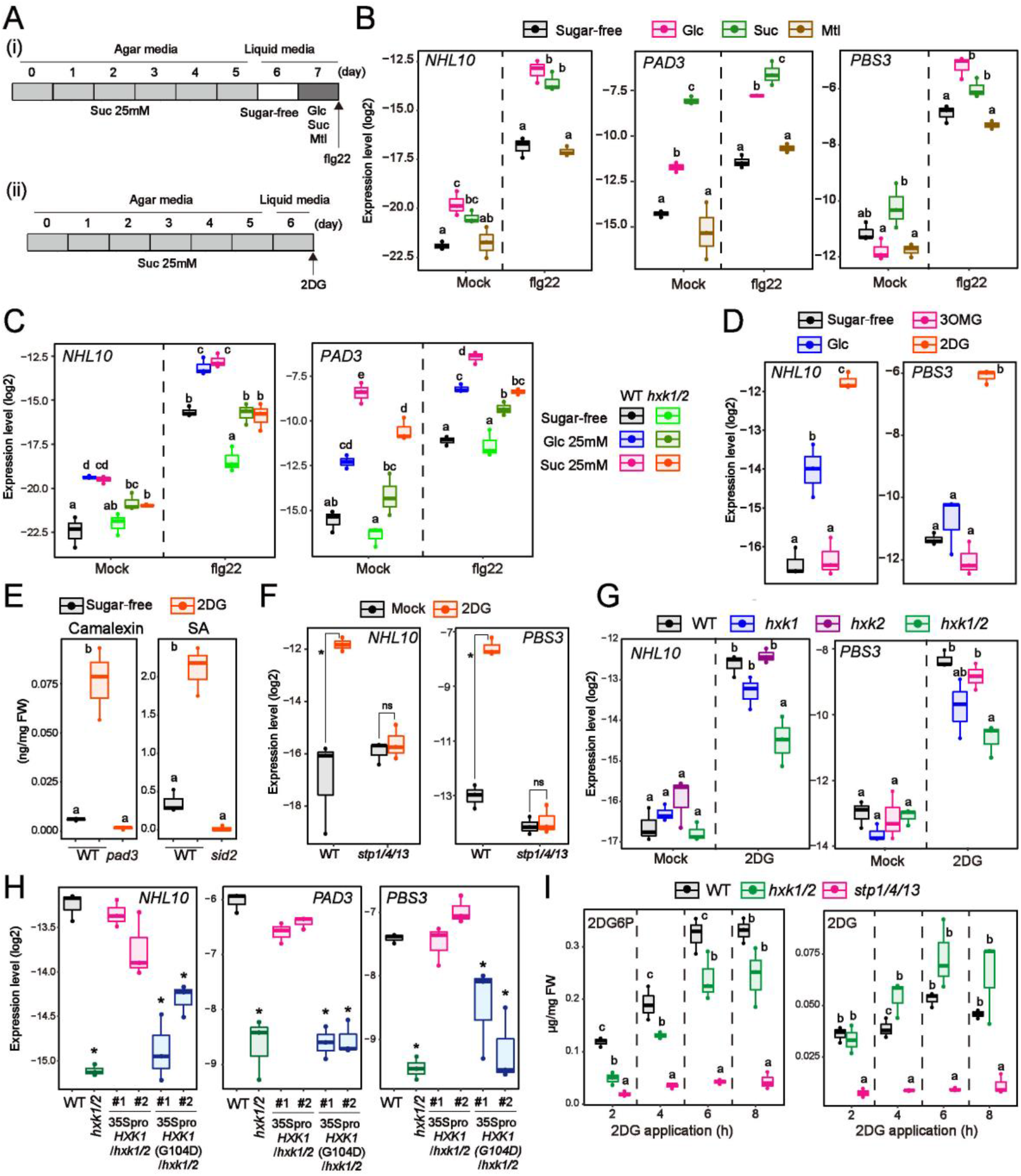
Sugar promotes plant defence signalling via HXK-mediated Glc phosphorylation. **A**, A schematic description of experimental procedures for flg22 application in the presence of glucose (Glc), sucrose (Suc), or mannitol (Mtl) (i) or 2DG treatment (ii). **B** and **C**, qRT-PCR analysis of defence-related gene expression in *Arabidopsis* seedlings exposed to mock or 1 µM flg22 for 8 h in the presence of 25 mM Glc, 25 mM Suc, or 25 mM Mtl (n = 3, biologically independent samples). **D**, qRT-PCR analysis of defence-related gene expression in *Arabidopsis* seedlings exposed to 25 mM Glc, 25 mM 3OMG or 0.5 mM 2DG for 24 h (n = 3, biologically independent samples). **E**, Quantification of camalexin and SA 24 h after 0.5 mM 2DG treatment (n = 3, biologically independent samples). FW indicates fresh weight. **F** and **G**, qRT-PCR analysis of defence-related gene expression in *Arabidopsis* seedlings exposed to mock or 0.5 mM 2DG for 8 h (n = 3, biologically independent samples). **H**, qRT-PCR analysis of defence-related gene expression in *Arabidopsis* seedlings exposed to 0.5 mM 2DG for 8 h (n = 3, biologically independent samples). **I**, Quantification of 2DG6P and 2DG after 0.5 mM 2DG treatment (n = 3, biologically independent samples). FW indicates fresh weight. Statistically significant differences among samples were determined using the two-tailed Welch’s t test (*P* <0.05) (E) or one-way ANOVA with multiple comparison tests (Tukey HSD) and are represented by asterisks or different letters (*P* <0.05). ns indicates non-significant.

Next, to explore the detailed mechanisms by which Glc activates defence signalling, we applied two non-metabolizable Glc analogues, 3-*O*-methyl-D-glucose (3OMG) and 2-deoxy-D-glucose (2DG), to plants for 24 h. Although both Glc analogues are imported into the cytoplasm via transporters (Boorer et al., 1994), only 2DG is phosphorylated by HXK (Jang and Sheen, 1994). *NHL10* expression was not affected by 3OMG but was significantly induced by 2DG (Figure 1D). Furthermore, RNA-seq analyses revealed that defence-related genes, including flg22-responsive genes, were overrepresented among 2DG-responsive genes (Figure S1A, S1B). Not only camalexin synthetic genes such as *PAD3*, but also SA-synthetic genes such as *AVRPPHB SUSCEPTIBLE 3* (*PBS3*) and *SALICYLIC ACID INDUCTION DEFICIENT 2* (*SID2*) were strongly induced in response to 2DG (Figure S1A, S3A). Consistently, camalexin and SA accumulated 24 h after 2DG application (Figure 1E). *PBS3* expression also responded to Glc and Suc. While Glc and Suc alone did not activate *PBS3* expression, they promoted flg22-triggered *PBS3* expression (Figure 1B).

2DG is a more potent inducer of many defence-related genes than Glc (Figure 1D, S1A). However, simultaneous application of excess Glc suppressed 2DG-induced expression to the levels under Glc conditions (Figure S3C), indicating competitive effects between 2DG and Glc on cellular activities such as transport and metabolism. Because 2DG is not further catabolized in glycolysis after HXK-mediated phosphorylation, 2DG-6-phosphate (2DG6P) accumulates following 2DG application (Figure S3D). However, we showed that cellular 2DG and 2DG6P decreased dramatically following simultaneous application of excess amounts of Glc (Figure S3D). Therefore, the amounts of cellular 2DG and/or 2DG6P appeared to correlate with the amplitude of defence signalling in response to 2DG. 2DG and 2DG6P act as inhibitors of HXK and phosphoglucose isomerase, respectively (Sols and Crane, 1954; Wick et al., 1957), thereby causing inhibition of glycolysis. We therefore deduced that glycolysis-derived energy is dispensable for 2DG-induced defence stimulation. In addition, we concluded that energy depletion caused by 2DG treatment does not lead to defence stimulation either, because the mitochondrial electron transport inhibitor antimycin A (AMA), the mitochondria uncoupler 2,4-dinitrophenol (DNP) and the photosynthesis inhibitor 3-(3,4-dichlorophenyl)-1,1-dimethylurea (DCMU) did not induce camalexin or SA (Figure S3E). The results described above suggested contributions of sugar transporters and HXK to 2DG-induced defence signalling. We focused on 8h after 2DG application, as at this time point 2DG responsiveness of most genes was observable (Figure S3A) and the secondary effect of SA accumulation was minimized because SA had not yet accumulated (Figure S3B). Of 14 *SUGAR TRANSPORT PROTEIN* (*STP*) genes that mainly transport monosaccharides in *Arabidopsis*, *STP1*, *STP4*, and *STP13* are the most abundantly expressed (Yamada et al., 2011). Therefore, we established *stp1 stp4 stp13* (*stp1/4/13*) triple mutants by introducing a 17 bp deletion in the *STP4* locus into *stp1/13* plants using CRISPR-Cas9 (Figure S4A-S4F). As expected, *stp1/4/13* plants exhibited a significant reduction in 2DG influx activity (Figure S4D). We found that *stp1/4/13* plants displayed no induction of *NHL10* or *PBS3* in response to 2DG (Figure 1F). Moreover, *hxk1/2* plants also showed reduced expression of *NHL10* and *PBS3* in response to 2DG (Figure 1G). Introduction of a *HXK1* cDNA complemented the reductions in 2DG-induced defence gene expression and 2DG6P production in *hxk1/2* plants (Figure 1H, S2H). On the other hand, introduction of *HXK1(G104D)* (Moore et al., 2003), which acts only as a Glc sensor and lacks catalytic activity, did not complement these phenotypes (Figure 1H, S2H). Thus, HXK-mediated 2DG phosphorylation is required for 2DG-induced defence signalling. However, we noticed that 2DG-induced *SID2* expression appeared to be restored by *HXK1(G104D)*, implying that HXK1’s sensor activity also contributes to 2DG-induced defence signalling (Figure S2G). We found that 2DG amounts increased in *hxk1/2* plants while 2DG6P decreased; *stp1/4/13* plants showed a reduction of both 2DG and 2DG6P (Figure 1I). Therefore, 2DG6P levels correlated with expression levels of defence-related genes in *hxk1/2* plants in response to 2DG. Together, these data suggested that 2DG6P, which HXK1/2 generate, primarily mediates defence signalling after 2DG application.

### Sugar stimulates the protein and phosphorylation levels of calcium-dependent protein kinase 5

Calcium-dependent protein kinases (CPKs/CDPKs), particularly CPK4/5/6/11 belonging to subgroup I, activate *NHL10* expression in response to flg22 (Boudsocq et al., 2010); moreover, CPK5/6 induce *PAD3* during fungal infection (Zhou et al., 2020). We found that loss of *CPK5/6* attenuated the induction of *NHL10* and *PAD3* in response to 2DG, while 2DG-induced expression of *PBS3* and *SID2* was enhanced in *cpk5/6* plants (Figure 2A). Consistently, accumulation of camalexin was almost abolished but that of SA was further increased in *cpk5/6* plants in response to 2DG (Figure 2B). CPK4/11 partially contributes to suppression of SA biosynthesis because *cpk4/5/6/11* plants showed higher SA accumulation than *cpk5/6* plants in response to 2DG, although 2DG-induced SA accumulation did not differ between WT and *cpk4/11* plants (Figure 2B). Introduction of a C-terminally GFP-fused *CPK5* genomic fragment containing its promoter region (gCPK5-GFP) fully complemented 2DG-induced gene expression in *cpk5/6* plants (Figure S5A). Therefore, CPK5/6, especially CPK5, affect 2DG-induced signalling by positively and negatively regulating the expression of *NHL10/PAD3* and *PBS3/SID2*, respectively, downstream of HXK1/2. Previously, plasma membrane-localised tobacco CDPKs, similar to *Arabidopsis* CPK5, were reported to be stimulated by application of Glc or Suc (Ohto and Nakamura, 1995). This is consistent with our data showing a relationship between sugar and CPKs. In addition, 2DG-induced *CAB1* attenuation was not affected by loss of CPK5/6, suggesting the presence of a CPK5/6-independent sugar signalling pathway(s) in *Arabidopsis* (Figure 2A).

**Figure 2.**
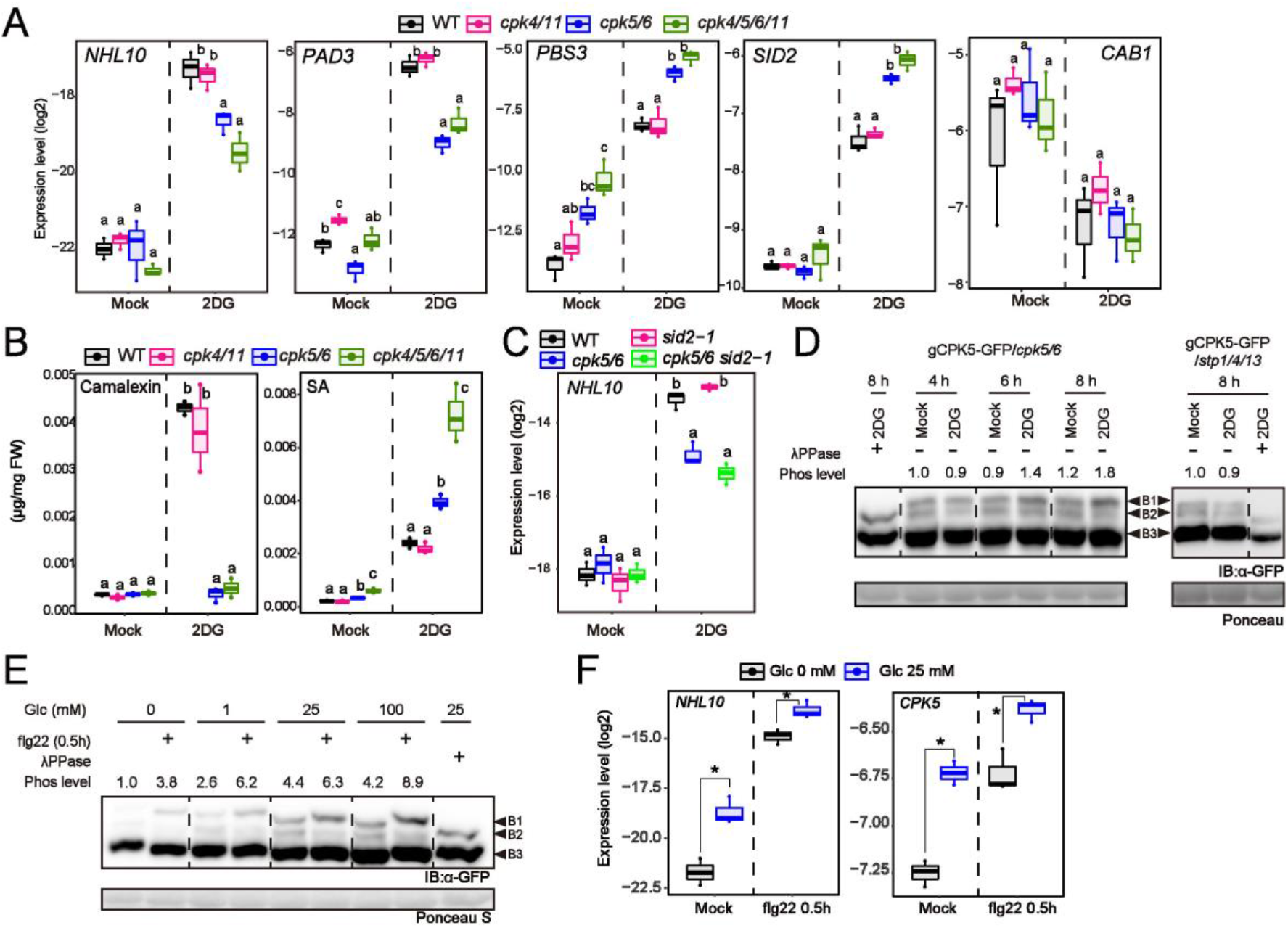
Clade A protein phosphatase C suppresses the activity of calcium-dependent protein kinase in a glucose-6-phosphoate-dependent manner. **A** and **C**, qRT-PCR analysis of defence-related gene expression in *Arabidopsis* seedlings exposed to mock or 0.5 mM 2DG for 8 h (n = 3, biologically independent samples). **B**, Quantification of camalexin and SA 24 h after mock or 0.5 mM 2DG treatment (n = 3, biologically independent samples). FW indicates fresh weight. **D** and **E**, Immunoblot analysis of CPK5-GFP in *Arabidopsis* seedlings after 0.5 mM 2DG application (D) or 1 µM flg22 application (E) (Mn^2+^-Phos-tag SDS-PAGE). Phospho levels were calculated as the ratios of intensities of B1 and B3. The phospho levels of mock 4h (D) and mock in the absence of Glc (E) were set to 1.0. IB denotes immunoblotting with anti-GFP antibodies. Ponceau S-stained membranes are shown below. **F**, qRT-PCR analysis of defence-related gene expression in *Arabidopsis* seedlings exposed to mock or flg22 for 0.5 h in the absence/presence of 25 mM Glc (n = 3, biologically independent samples). Individual data points are plotted. In box plots, centre lines represent the medians, box edges delimit lower and upper quartiles and whiskers show the highest and lowest data points. Experiments in this figure were repeated at least three times with similar trends. Statistically significant differences among samples were determined using the two-tailed Welch’s t test (*P* <0.05) (A, F) or one-way ANOVA with multiple comparison tests (Tukey HSD) and are represented by asterisks or different letters (*P* <0.05). ns indicates non-significant.

*NHL10* expression was similar in *cpk5/6* and *cpk5/6/sid2* plants (Figure 2C, S5B), indicating that enhanced SA levels in *cpk5/6* plants did not reduce 2DG-induced *NHL10* expression. CPK5/6 phosphorylate the transcription factors WRKY33 (group I) and WRKY8/28/48 (group IIc), transducing signals to downstream pathways (Gao et al., 2013; Zhou et al., 2020). Expression of *NHL10* and *PAD3* in response to 2DG was reduced in *wrky33* plants, but not *wrky8* plants, (Figure S5C), suggesting that WRKY33 acts as a downstream factor of CPK5/6 in 2DG-induced signalling.

We next asked if 2DG application affects CPK5 status. An immunoblot showed three bands of CPK5-GFP (Figure 2D). Of these, the uppermost band (B1) completely disappeared after treatment with lambda protein phosphatase (λPPase) (Figure 2D), indicating that B1 represents the major phosphorylated form of CPK5-GFP. CPKs are rapidly activated upon PAMP perception following Ca^2+^ influx (Boudsocq et al., 2010). We found that flg22 application increased the intensity of B1 and reduced that of the middle band (B2), but did not affect the lowest band (B3) of CPK5-GFP after transient elevation of the cytosolic Ca^2+^ concentration (Figure S6A). These data indicated that the B2 form of CPK5-GFP becomes phosphorylated and shifts to the B1 form in response to flg22. We found that the B3 form is an alternative splicing variant because it was only faintly observable in plants expressing CPK5(cDNA)-GFP from the constitutive 35S promoter (Figure S6B). We evaluated the dynamics of CPK5 phosphorylation by calculating the B1/B3 ratio of CPK5-GFP. The B2 intensity was sometimes too weak to use for normalization, so instead we used the stable signal of B3 as an indicator of CPK5 protein abundance. The CPK5 phosphorylation level (B1/B3 ratio) indeed increased under 2DG treatment (Figure 2D), but there was no such increase in *stp1/4/13* plants, indicating that 2DG uptake leads to CPK5 phosphorylation.

Next, we investigated the phosphorylation status of CPK5-GFP in the presence of Glc. Consistent with the 2DG experiments, phosphorylation levels of CPK5-GFP increased under higher Glc conditions (Figure 2E). The protein abundance of CPK5-GFP was elevated in a Glc-dependent manner (Figure 2E). We found that *CPK5* expression was induced by Glc application (Figure 2F), likely contributing to an increase in CPK5 protein amounts. On the other hand, flg22-triggered CPK5 phosphorylation occurred even in the absence of Glc (Figure 2E), suggesting that Glc is dispensable for flg22-triggered CPK5 phosphorylation. However, a Glc-dependent increase in basal phosphorylation levels supported higher flg22-triggered phosphorylation levels of CPK5-GFP (Figure 2E). Indeed, *NHL10* expression 0.5 h after flg22 application appeared to correlate with protein and phosphorylation levels of CPK5-GFP (Figure 2F). Taken together, these data suggested that Glc primes CPK5 activity for defence gene expression by increasing its protein and phosphorylation levels.

### Dephosphorylation of CPK5 by clade A protein phosphatase 2Cs is suppressed by glucose-6-phosphate

Despite inducing CPK5 phosphorylation, 2DG treatment did not trigger discernible elevation of [Ca^2+^]cyt (Figure 3A). This prompted us to explore Ca^2+^-independent CPK5 regulatory mechanisms. CPK6 and CPK11 were previously reported to bind to clade A PP2Cs (Brandt et al., 2015; Lynch et al., 2012). We asked if CPK5 also interacts with clade A PP2Cs in yeast two-hybrid and split-luciferase assays. Of clade A PP2Cs we tested, we found that ABI1 and PP2CA strongly, and HAI1 weakly, bound to CPK5, whereas HAB1 did not (Figure 3B, S6C). On the other hand, AP2C1 (clade B) and PP2C6-6 (clade E) did not bind to CPK5 (Figure S6D), suggesting that CPK5 interacted specifically with clade A PP2Cs. Consistent with the protein interaction data, *in vitro* CPK5 phosphorylation levels, detected by anti-phosphothreonine (anti-pThr) antibody, were massively reduced by ABI1 and PP2CA, but not by HAB1 (Figure S6E). Furthermore, CPK5 *trans-*phosphorylation activity on an N-terminal fragment of RBOHD (RBOHD-NT) (Kadota et al., 2014) and a WRKY33 wbox1 fragment (Zhou et al., 2020) was weakened in the presence of ABI1 (Figure 3C, S6F). Consistent with previous reports (Geiger et al., 2010; Lynch et al., 2012), *in vitro trans-*phosphorylation activity of the protein kinase OPEN STOMATA 1 (OST1) was also suppressed in the presence of ABI1 (Figure S6G). Importantly, when we added ABI1 to the reaction after stopping CPK5 activity using the kinase inhibitor staurosporine, the phosphorylation levels of CPK5 substrates were not reduced (Figure 3C, S6F). Therefore, we can exclude the possibility that ABI1 directly dephosphorylated CPK5 substrates. Interestingly, this ABI1 effect on CPK5 activity was alleviated in the presence of Ca^2+^ (Figure 3C), indicating that Ca^2+^-activated CPK5 can overcome ABI1-mediated suppression. We inferred from these data that clade A PP2Cs suppress CPK5 activity, especially before PAMP-triggered Ca^2+^ influx.

**Figure 3.**
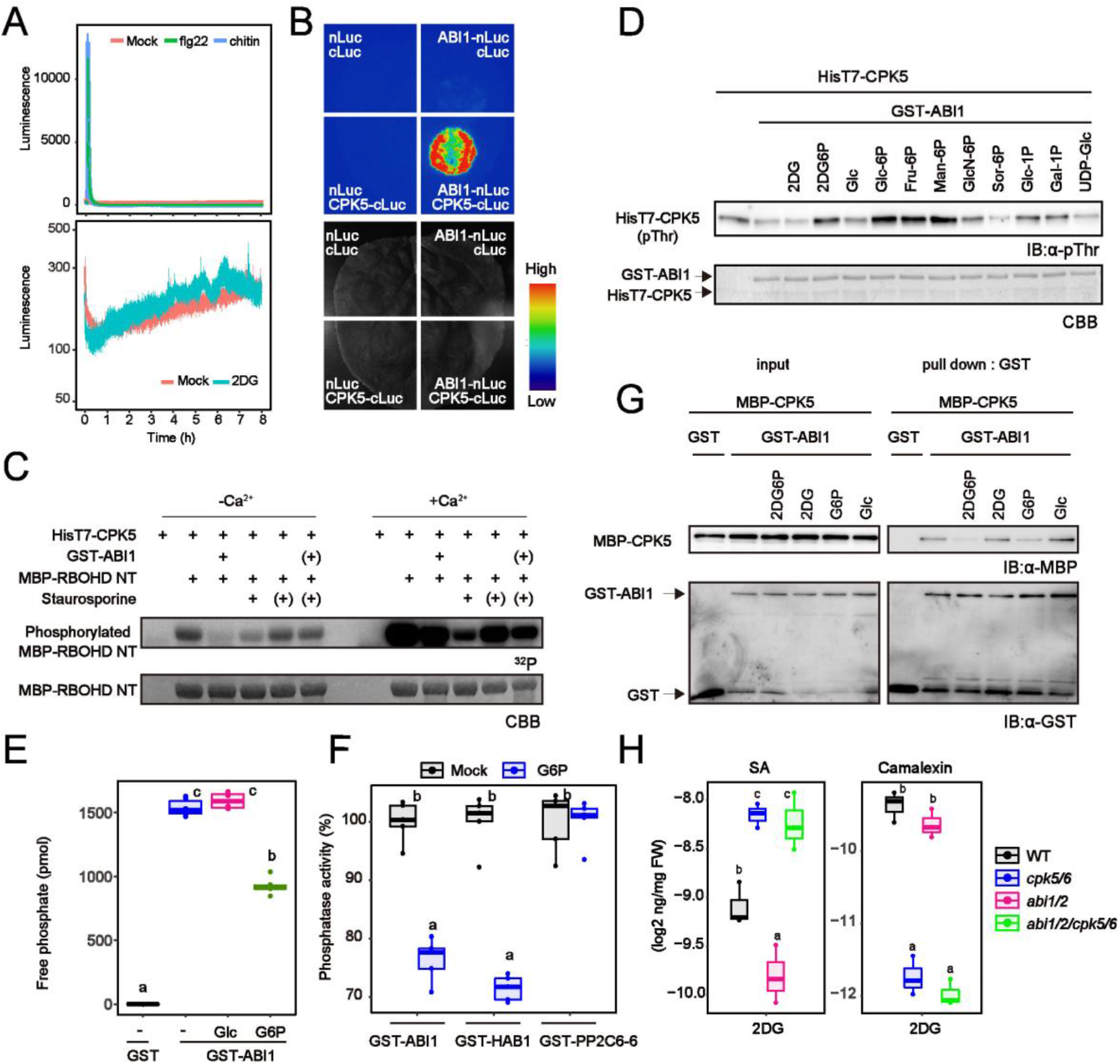
Glucose-6-phosphate suppresses dephosphorylation of CPK5 by clade A protein phosphatase 2Cs. **A**, Cytosolic Ca^2+^ concentrations were measured via aequorin-mediated luminescence in response to PAMPs (left) or 2DG (right) (mean ± SE, n = 8, biologically independent samples). **B,** Split-luciferase assay for protein interactions between ABI1 and CPK5 in *N. benthamiana* leaves. **C**, Autoradiograph of RBPHD-NT phosphorylation by CPK5 *in vitro* for 30 min with [γ-^32^P] ATP (upper images). Parentheses indicate addition 15 min after the reaction was started. CBB-stained gels are shown below. **D**, ABI1 dephosphorylate CPK5 *in vitro*. Phosphorylation of CPK5 was detected by anti-phopsho-threonine antibody. 10 mM sugars were applied. IB denotes immunoblotting. Fru-6P, Man-6P, GlcN-6P, Sor-6P, Glc-1P, Gal1P, and UDP-Glc indicate fructose-6-phosphate, mannose-6-phosphate, glucosamine-6-phosphate, sorbitol-6-phosphate, glucose-1-phosphate, galactose-6-phosphate, and UDP-glucose, respectively. **E** and **F**, *in vitro* phosphatase activity of ABI1. 10 mM sugars were applied. G *in vitro* protein interaction between CPK5 and ABI1. 10 mM sugars were applied. IB denotes immunoblotting with anti-MBP or anti-GST antibodies. **H**, Quantification of camalexin and SA 24 h after mock or 0.5 mM 2DG treatment (n = 3, biologically independent samples). FW indicates fresh weight. Statistically significant differences among samples were determined using one-way ANOVA with multiple comparison tests (Tukey HSD) and are represented by different letters (*P* <0.05). ns indicates non-significant.

We next asked if sugar affects ABI1-mediated CPK5 regulation. Because we found that 2DG6P is involved in 2DG-induced defence signalling, we investigated whether 2DG6P and G6P which is the natural counterpart of 2DG6P affect ABI1-mediated CPK5 dephosphorylation. We demonstrated here that application of 2DG6P and G6P but not 2DG or Glc reduced ABI1-mediated CPK5 dephosphorylation *in vitro* (Figure 3D). We found that G6P suppressed the *in vitro* protein phosphatase activities of ABI1 and HAB1 (Figure 3E, 3F, S6I), while the activity of PP2C6-6 (clade E) was not affected (Figure 3F). In addition, λPPase-mediated CPK5 dephosphorylation was not reduced by G6P (Figure S6H), implying that G6P specifically influences the activity of clade A PP2Cs. Furthermore, we revealed that the application of 2DG6P and G6P, but not 2DG or Glc, dissociated protein interactions between CPK5 and ABI1 (Figure 3G). Together, these results suggest that both 2DG6P and G6P restored CPK5 phosphorylation levels by suppressing the activity of clade A PP2Cs and dissociating clade A PP2Cs from CPK5.

Other hexose-phosphates including hexose-1-phosphates, but not the sugar nucleotide UDP-Glc, also inhibited ABI1-mediated CPK5 dephosphorylation, whereas Glc-1-phosphate had a weaker effect than G6P (Figure 3D, S6J). In contrast, sorbitol-6-phosphate did not affect ABI1 activity (Figure 3D), suggesting that the phosphate group did not directly inhibit ABI1 activity. G6P concentration was previously reported to be around 3-5 mM in the cytoplasm of plant cells (Leidreiter et al., 1995). G6P at this concentration may not be enough to suppress PP2C activity; G6P above 10 mM inhibited ABI1 activity *in vitro* (Figure S6I). We speculate that combinations of G6P and other sugar phosphates may lead to suppression of PP2C activity in plant cells under sugar-sufficient conditions.

We next investigated the relationship between CPK5/6 and ABI1/2 in plants. While camalexin synthesis in response to 2DG was similar in WT and *abi1/2* plants, 2DG-induced SA was reduced in *abi1/2* plants, compared with WT plants (Figure 3H). On the other hand, SA accumulation in response to 2DG was elevated in *abi1/2/cpk5/6* plants to a similar extent as in *cpk5/6* plants (Figure 3H), implying that enhanced CPK5/6 activity caused a reduction in 2DG-induced SA synthesis in *abi1/2* plants.

### Sugar influx contributes to pattern-triggered immunity

We have reported that elevated sugar influx activity in response to flg22 interrupts pathogens’ sugar acquisition (Yamada et al., 2016). Given that, while apoplasmic sugar amounts decrease, cytoplasmic sugar should concomitantly increase in the cytoplasm during flg22-triggered immunity. Indeed, we found that G6P amounts were elevated in response to flg22 under 25 mM Suc conditions, dependent on the activation of sugar transporter STPs (Figure 4A, 4B). Flg22-triggered expression of defence-related genes decreased in *hxk1/2* plants in the presence of 25 mM Glc or Suc (Figure 1C). Flg22-triggered synthesis of SA and camalexin was also reduced in *hxk1/2* plants under these sugar conditions (Figure S7A), suggesting that G6P, which HXK1/2 generate, affects flg22-triggered defence responses. Together, we hypothesized that STP-mediated sugar influx contributes to flg22-triggerd signalling. We showed that sugar influx activity was enhanced 8 and 24 h, but not 3h, after flg22 application (Figure S7B). Therefore, to focus on the contribution of STP-mediated sugar influx to flg22-triggered signalling during these later time points, we treated plants with flg22 under 25 mM Suc conditions. Whereas possible differences in cellular sugar amounts before PAMP recognition would cause different CPK5 priming effects between WT and *stp1/4/13* plants, this effect can be minimized under sugar-sufficient conditions. In addition, Suc influx is likely unaffected when the monosaccharide transporters STPs are absent. Notably, RNA-seq analyses revealed that transcriptional profiles in response to flg22 in the presence of 25 mM Suc were substantially altered in *stp1/4/13* plants compared with WT plants (Figure 4C, S7C). As expected, these differences in transcriptional profiles increased at the later stage of flg22-triggered signalling (Figure 4C), correlating with the timing of enhancement of sugar uptake activity in response to flg22 (Figure S7B). Of particular interest was the observation that STP-dependent genes largely overlapped with 2DG-responsive genes during flg22-triggered defence (Figure 4C). This suggests that sugar signalling activated by STP-mediated sugar influx stimulates defence signalling in response to flg22. In addition, *stp1/4/13* plants showed higher susceptibility to the less virulent strain of the pathogenic bacterium *Pseudomonas syringae* pv. *tomato* (*Pst*) DC3000, *Pst* DC3000 (*ΔavrPtoΔavrPtoB*) (Figure 4F).

**Figure 4.**
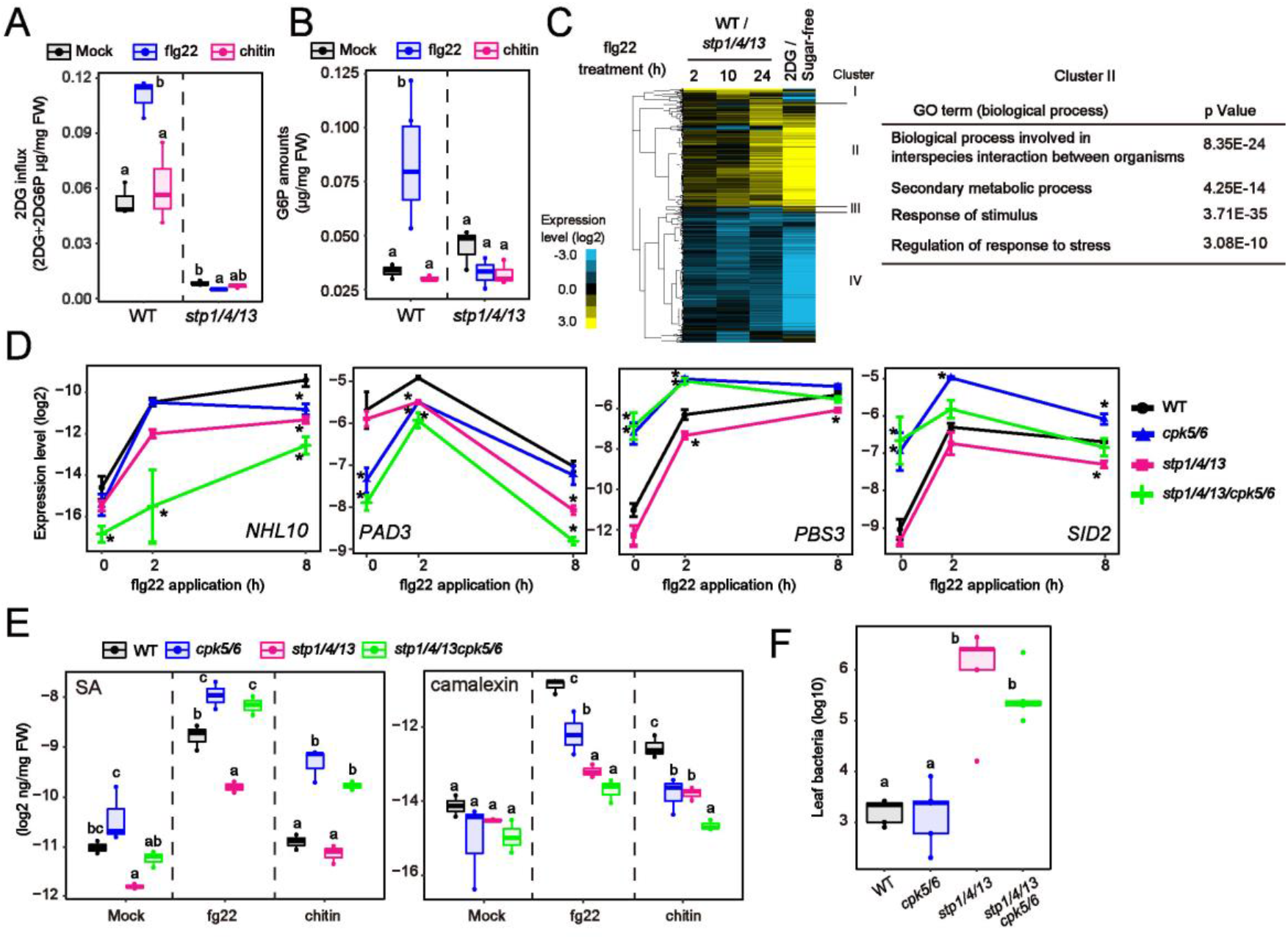
Sugar influx contributes to pattern-triggered signalling. **A**, Glucose influx activity in *Arabidopsis* seedlings 8 h after 1 µM flg22 treatment (n = 3, biologically independent samples). Glucose influx activity was measured by calculating the mixed amounts of 2DG and 2DG6P 1h after 0.5 mM 2DG application. **B**, Quantification of G6P in *Arabidopsis* seedlings 8 h after 1 µM flg22 treatment under 25 mM Suc conditions (n = 3, biologically independent samples). FW indicates fresh weight. **C**, Heat maps displaying expression patterns of genes that show significant expression changes in *stp1/4/13* plants, relative to WT plants, in response to 1 µM flg22 in the presence of 25 mM Suc (q < 0.01 and |log_2_FC| > 1). The log_2_ fold changes relative to WT were subjected to hierarchical clustering analysis (left). Overrepresented GO terms in cluster II are presented (right). **D**, qRT-PCR analysis of defence-related gene expression in *Arabidopsis* seedlings exposed to 1 µM flg22 under 25 mM Suc conditions (n = 3, biologically independent samples). **E**, Quantification of SA and camalexin in *Arabidopsis* seedlings 8 h after 1 µM flg22 treatment under 25 mM Suc conditions (n = 3, biologically independent samples). FW indicates fresh weight. **F**, Growth of *Pst* DC3000(*ΔavrPtoΔavrPtoB*) spray-inoculated onto rosette leaves of 4-week-old *Arabidopsis*. Bacterial titres were determined 3 days post-inoculation (n = 5, biologically independent samples). Statistically significant differences among samples were determined using one-way ANOVA with multiple comparison tests (Tukey HSD or Dunnett’s test between WT and other genotypes at each time point (D)) and are represented by asterisks or different letters (*P* <0.05).

We found that, in the presence of 25 mM Suc, loss of *CPK5/6* reduced flg22-triggered expression of *NHL10* and *PAD3* to a similar extent as loss of *STP1/4/13* (Figure 4D). In addition, flg22-triggered camalexin synthesis was also reduced in *cpk5/6* and *stp1/4/13* plants. We expected that STP-mediated sugar influx would contribute to flg22-triggered defence gene expression by enhancing CPK5/6 activity. However, CPK5 phosphorylation levels were not altered between WT and *stp1/4/13* plants 8h after flg22 application (Figure S7D), at which time G6P accumulates via sugar influx (Figure 4A, 4B). We showed that Ca^2+^-activated CPK5 overcomes ABI1-mediated dephosphorylation (Figure 3C). Although flg22-triggered Ca^2+^-influx has already returned to baseline by 8h after flg22 application (Figure 3A), CPK5, once activated by Ca^2+^, may not be affected by G6P for some time. Next, to dissect the relationship between CPK5/6-mediated and STP-mediated signalling during PTI, we established *stp1/4/13/cpk5/6* quintuple mutants (Figure S8A). These *stp1/4/13/cpk5/6 p*lants showed almost no response to 2DG, like *stp1/4/13* plants (Figure S8B). Thus, *stp1/4/13* mutations are epistatic to *cpk5/6* mutations with respect to 2DG response because 2DG influx is essential to trigger 2DG-induced signalling. On the other hand, in response to flg22, *stp1/4/13/cpk5/6* plants showed an additive phenotype relative to *cpk5/6* and *stp1/4/13* plants. *stp1/4/13/cpk5/6* plants exhibited a further reduction in flg22-triggered expression of *NHL10* and *PAD3*, compared with *cpk5/6* and *stp1/4/13* plants (Figure 4D). These additive effects suggested that STP-mediated sugar influx stimulates CPK5/6-independent signalling, which promotes flg22-triggered expression of *NHL10* and *PAD3* together with Ca^2+^-activated CPK5/6-mediated signalling.

Expression of *PBS3* and *SID2* in response to flg22 were also reduced in *stp1/4/13* plants (Figure 4E). Consistently, flg22-induced SA amounts were less induced in *stp1/4/13* plants (Figure 4D), indicating that sugar influx activates SA biosynthesis pathway in response to flg22. On the other hand, we found that SA-synthetic gene expression and SA accumulation were highly induced in *cpk5/6* plants in response to flg22 as well as 2DG (Figure 4D, 4E), although CPK5 was previously reported to promote SA synthesis during effector-triggered immunity (Gao et al., 2013; Liu et al., 2017). We confirmed that enhanced SA amounts in *cpk5/6* plants were also detected in mature leaves after flg22 application or inoculation of *Pst* DC3000 (Figure S9A). In addition, although *cpk5/6* plants were less susceptible to spray-inoculated *Pst* DC3000 than WT plants, an SA-deficient *sid2* mutation conferred higher susceptibility to *Pst* DC3000 to *cpk5/6* plants (Figure S9B), suggesting that high SA levels elevated anti-bacterial resistance in *cpk5/6* plants under our experimental conditions. Interestingly, we demonstrated that *cpk5/6* mutations were epistatic to *stp1/4/13* mutations with respect to flg22-triggered SA synthesis. In response to flg22, *stp1/4/13/cpk5/6* plants showed high SA accumulation similar to *cpk5/6* plants, whereas SA levels were reduced in *stp1/4/13* plants (Figure 4E). These results revealed that CPK5/6-mediated SA suppression becomes dominant in *stp1/4/13* plants, and suggested that STP-mediated sugar signalling promote SA synthesis by attenuating this CPK5/6 effect in WT plants. However, we guess that STP-mediated sugar influx does not lead to inhibition of CPK5/6 activity, because flg22-triggered SA synthesis was not altered between WT and *cpk5/6* plants under sugar-deficient conditions (Figure S9C), which indicates that loss of CPK5/6 activity was not a direct trigger for SA synthesis. Therefore, we concluded that STP-mediated sugar influx enhances SA-synthetic signalling which antagonistically acts on CPK5/6-mediated SA suppression during flg22-triggered immunity. On the other hand, elevated susceptibility to *Pst* DC3000 (*ΔavrPtoΔavrPtoB*) in *stp1/4/13* plants was not significantly reduced in *stp1/4/13/cpk5/6* plants despite higher SA accumulation, whereas it decreased mildly (Figure 4F). Because an increase in apoplasmic sugar amounts by loss of STPs elevates bacterial virulence (Yamada et al., 2016), it could cancel SA-mediated immunity from loss of CPK5/6.

We found that, unlike flg22, application of the fungal cell wall component chitin did not stimulate sugar influx activity or increase G6P amounts (Figure 4A, 4B). *STP13*, a gene responsible for flg22-induced sugar influx activity (Yamada et al., 2016), was less induced in response to chitin than in response to flg22 (Figure S10A, S10B). Moreover, CERK1, the co-receptor for chitin receptors, did not phosphorylate the STP13 T485 residue (Yamada et al., 2016) which is the important residue for elevation of sugar influx activity (Figure S10C), although BAK1, the co-receptor for the flagellin receptor, did *in vitro* (Yamada et al., 2016). Therefore, we wondered if sugar influx activity does not contribute to chitin-triggered signalling. However, chitin-triggered *PAD3* expression and camalexin synthesis were reduced in *stp1/4/13* plants (Figure 4E, 5A), indicating that basal sugar influx activity mediated by STPs is required for chitin-triggered signalling. On the other hand, chitin-triggered *NHL10* expression was not reduced, while flg22-triggered expression was reduced, in *stp1/4/13* plants (Figure 5A), suggesting that the contribution of STP-mediated sugar influx to chitin-triggered signalling is lower than to flg22-triggered signalling. STP-mediated sugar influx led to activation of CPK5-independent signalling pathways in response to chitin as well as flg22. *PAD3* expression and camalexin accumulation were further reduced in *stp1/4/13/cpk5/6* plants in response to chitin, compared to *cpk5/6* and *stp1/4/13* plants (Figure 4E, 5A). Furthermore, anti-fungal defence against the necrotrophic fungus *Alternaria brassicicola* was massively reduced in *stp1/4/13/cpk5/6* plants (Figure 5B). Taken together, these data show that STP-mediated sugar influx contributes to defence against both bacterial and fungal pathogens, although it differentially affects these PAMP receptor signalling processes.

**Figure 5.**
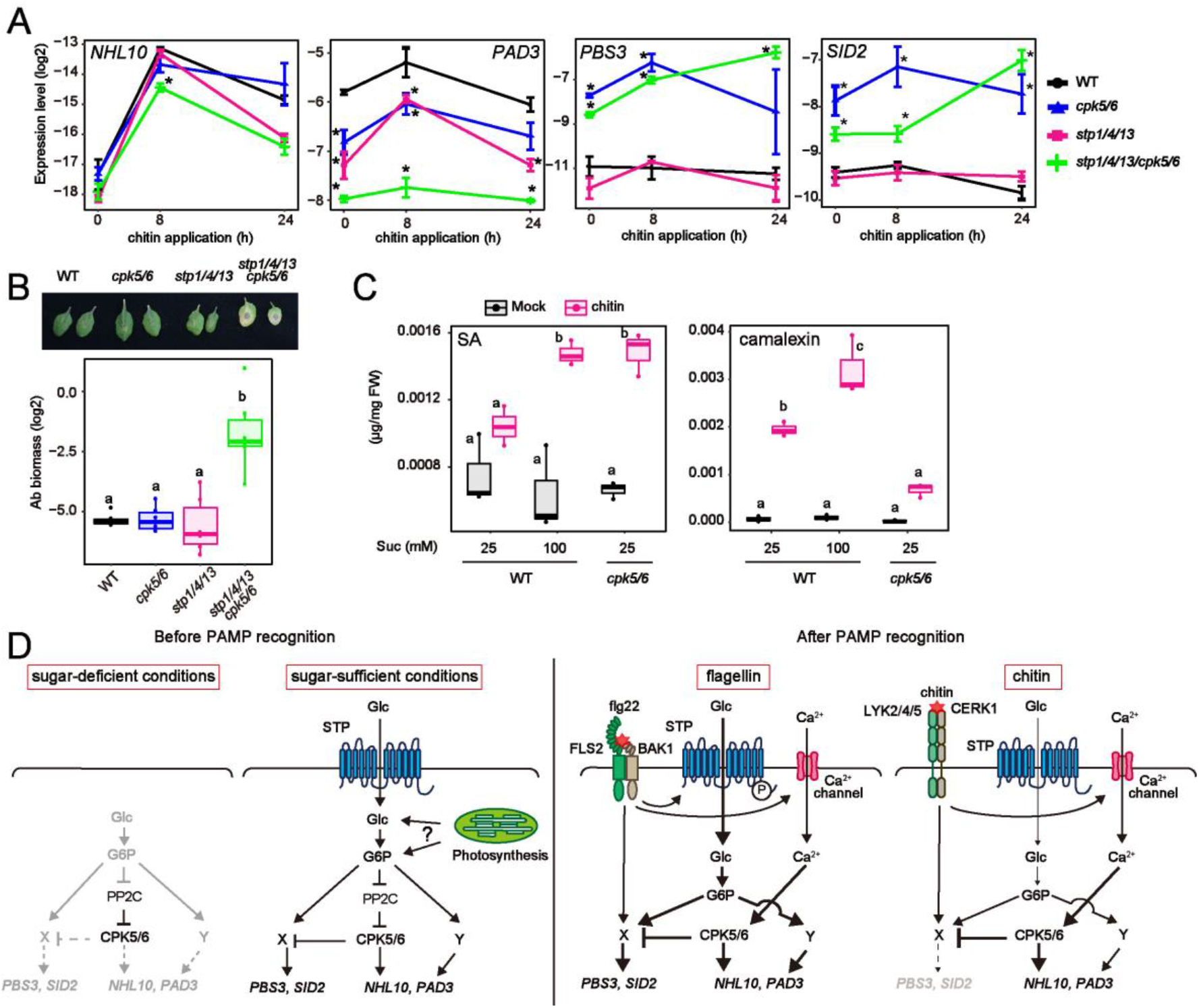
Cellular sugar level is key to co-ordinating defence outputs during PTI. **A**, qRT-PCR analysis of defence-related gene expression in *Arabidopsis* seedlings exposed to chitin under 25 mM Suc conditions (n = 3, biologically independent samples). **B**, Biomass of *A. brassicicola* on detached rosette leaves of 4-week-old *Arabidopsis* after 5 days (n = 4, biologically independent samples). **C**, Quantification of SA and camalexin in *Arabidopsis* seedlings 8 h after chitin treatment under 25 mM or 100 mM Suc conditions (n = 3, biologically independent samples). FW indicates fresh weight. **D**, A model proposed in this study. An increase in cellular G6P, possibly through sugar influx and photosynthesis, results in suppression of clade A PP2Cs to prime CPK5/6. During PTI, sugar influx triggers CPK5-independent signalling. One pathway (Y) additively activates *NHL10*/*PAD3* with CPK5/6. The other pathway (X) is antagonistic to CPK5/6 which suppress *PBS3*/*SID2*. Sugar influx activity is enhanced in response to flg22, leading to SA accumulation that promotes anti-bacterial defence. On the other hand, chitin recognition only promotes camalexin synthesis, because sugar influx activity is not affected. Statistically significant differences among samples were determined using one-way ANOVA with multiple comparison tests (Tukey HSD or Dunnett’s test between WT and other genotypes at each time point (A)) and are represented by asterisks or different letters (*P* <0.05).

### Cellular sugar amounts determine SA synthesis during pattern-triggered immunity

We found that chitin application did not induce SA-synthetic gene expression or SA synthesis under our experimental conditions (25 mM Suc) (Figure 5A). Previous studies also reported that chitin treatment did not induce expression of the SA marker gene *PR1* (Gust et al., 2007). However, chitin treatment induced SA accumulation in *cpk5/6* plants (Figure 5C). This result suggests that chitin signalling can stimulate SA-synthetic signalling, but is too weak to overcome CPK5/6-mediated suppression. We noticed that it resembles flg22-triggered SA accumulation patterns in *stp1/4/13* and *stp1/4/13/cpk5/6* plants. Therefore, unincreased cytoplasmic sugar levels (Figure 4B), due to unenhanced sugar influx activity (Figure 4A), may cause undetectable chitin-triggered SA synthesis. To investigate this hypothesis, we treated plants with chitin under higher sugar conditions (100 mM Suc) to elevate basal cellular sugar amounts. Notably, chitin-triggered SA accumulation was observable even in WT plants under these conditions (Figure 5C). Taken together, these data suggested that cellular sugar levels are key factors for SA synthesis during PTI.

## Discussion

In this study, we provide evidence that cellular sugar level is a critical factor regulating defence outputs. We found that G6P, not Glc, contributes to defence activation in *Arabidopsis*. Likewise, G6P has been reported to stimulate sugar signalling in animals, for instance via promoting the nuclear localization of the transcription factor MondoA (Stoltzman et al., 2008). Because Glc is rapidly phosphorylated by HXK in the cytoplasm, Glc amounts are limited in the cytoplasm; Glc is mainly compartmentalized into the vacuole in plant cells (Leidreiter et al., 1995). Therefore, cytoplasmic G6P sensory mechanisms may have evolved in various organisms. However, the mode of action of these G6P sensors has been elusive. We propose a mechanism by which G6P suppresses the activity of clade A PP2Cs and dissociates the PP2Cs from CPK5, leading to an increase in CPK5 phosphorylation levels. This contributes to defence priming for stronger PTI responses.

After recognition of PAMPs, STP-mediated sugar influx activates CPK5-independent signalling. We showed that at least two CPK5-independent pathways are stimulated by sugar influx during PTI. One stimulates *NHL10* and *PAD3*, like the CPK5/6 pathway (Figure 5D). The other acts antagonistically on the CPK5/6 pathway which represses *PBS3* and *SID2* (Figure 5D). The later signal pathway is required to synthesize SA during PTI. The sugar sensor SnRK1 becomes active, prioritizing the catabolic rather than the anabolic pathway under sugar-starved conditions. It was reported that *SnRK1* overexpression led to suppression of Glc-induced *PR1* (Jossier et al., 2009). Because G6P is a candidate molecule for suppression of SnRK1 activity (Toroser et al., 2000), enhanced G6P levels via STPs could inhibit SnRK1 activity for SA synthesis in response to flg22.

SA signalling is critical for anti-bacterial defence although it can have negative effects on anti-fungal defence, especially against necrotrophic fungal pathogens (Flors et al., 2008). Our study demonstrated that flg22-triggered signalling forces sugar transporters to elevate cellular sugar amounts, activating SA synthesis. In contrast, basal sugar amounts determined SA synthesis during chitin-triggered signalling. Chitin signalling did increase SA synthesis only in the presence of high levels of sugar. We inferred from these data that flg22 and chitin receptor signalling differentially affect sugar influx activity, promoting appropriate defence outputs against pathogens with different sensitivities to SA-mediated defence. However, sugar influx activity was reported to be enhanced during infection by necrotrophic fungal pathogens (Veillet et al., 2017), suggesting that fungal PAMPs other than chitin could activate sugar influx activity. As the synthesis of not only SA but also camalexin in response to chitin was elevated under higher Suc conditions (Figure 5C), sugar influx also contributes to increasing the amplitude of anti-fungal defence.

We showed qualitative differences between flg22-triggered and chitin-triggered defence outputs in a manner that depends on cellular sugar amounts. Although recognition of each PAMP immediately induces similar sets of defence-related genes (Bjornson et al., 2021), transcriptional profiles triggered by each PAMP during later phases were reported to be different (Kim et al., 2014). We expected that sugar influx might cause these differences during the later PTI phases. In fact, the contribution of STPs became larger during the late phase of flg22-triggered signalling, apparently correlating with the timing of enhancement of STP-mediated sugar influx activity (Figure S7B). Because nutrient competition occurs at the interface of host-pathogen interactions, we previously described sugar uptake by STPs as contributing to interruption of pathogen sugar acquisition (Yamada et al., 2016). We now highlight it again as an important plant defence system that not only suppresses the pathogen but also co-ordinates the plant defence. Altogether, we provide a comprehensive view of how sugar contributes to defence signalling in plants. Because defence signalling is energy-intensive, estimating sugar availability by monitoring cellular G6P amounts may allow fine-tuning of defence outputs. Furthermore, we propose that host metabolic conditions are involved in activating appropriate plant defence outputs. Environmental cues such as light, humidity, and temperature dynamically influence cellular sugar levels (Gibon et al., 2004; Maruyama et al., 2009). These environments were also reported to affect plant defence signalling. For example, dark-induced sugar-starvation elevated susceptibility to pathogens (Engelsdorf et al., 2013). In addition, cold and drought conditions, which increase cellular sugar amounts in plants, induced *PR1* expression (Kim et al., 2017; Liu et al., 2013). It will be interesting to analyse whether sugar plays a role in molecular integration of environmental conditions into defence signalling.

## Materials and Methods

### Plant materials and growth conditions

The Columbia-0 (Col-0) ecotype of Arabidopsis thaliana was used as wild type. Col-0 mutants stp1 (SALK_048848)^9^, stp13 (SALK_045494)^11^, cpk4 (SALK_000685)^35^, cpk5 (SAIL_657_C06)^15^, cpk6 (SALK_025460)^15^, cpk11 (SALK_054495)^12^, wrky33-1 (SALK_006603)^36^, wrky8-1 (SALK_107668C)^15^, hxk1 (WiscDsLoxHs044_02E), hxk2^37^.(WiscDsLox289_292O3), sid2-137 and pad3-138 were used. To establish the stp1 stp4 stp13 mutant, a 17 bp deletion in the STP4 locus was introduced into stp1 stp13 plants using CRISPR-Cas9. To establish stp4, stp1 stp4, and stp4 stp13 plants, stp1 stp4 stp13 plants were crossed with Col-0 plants, and the desired genotypes were detected from the F2 progeny. pMAQ239 plants were used to measure [Ca2+]cyt. Plants were grown on soil or 0.5× Murashige and Skoog (MS) agar medium (0.5× MS salt (Wako), 0.5× Gamborg’s vitamins (Sigma), 25 mM sucrose, 0.5 g/l 2-Morpholinoethanesulfonic acid monohydrate (MES) (pH 5.7), 0.8% agar) at 22 °C under 10 h light / 14 h dark or 16 h light / 8 h dark conditions, respectively.

### Plasmid construction

The 35S GFP NosT fragment was amplified from pRI 35S-GFP^20^, and inserted into the HindIII/EcoRI sites of pCAMBIA1300. This plasmid was named pCAMBIA 35S-GFP. To establish transgenic Arabidopsis, a genomic fragment including the upstream and coding regions of CPK5 or STP13 was inserted into the HindIII/SalI sites of pCAMBIA 35S-GFP, or a CPK5 cDNA fragment was inserted into the XbaI/SalI site of pCAMBIA 35S-GFP. A cDNA fragment of HXK1 or HXK1(G104D) was inserted into the XbaI/SalI sites of pCAMBIA 35S-GFP. For obtaining recombinant proteins, cDNA fragments of CPK5 or OST1 were inserted into the BamHI/HindIII sites of pET-28a(+). cDNA fragments of ABI1, HAB1, PP2C6-6, ABF1, HXK2, or HXK2ΔC13 were inserted into the BamHI/SalI sites of pGEX-6P-1. cDNA fragments of PP2CA, N-terminal truncated RBOHD, and WRKY33 wbox1 were inserted into the EcoRI/SalI site of pMAL-c2. For yeast two-hybrid assays, cDNA fragments of ABI1, HAB1, PP2CA, or HAI1 were inserted into the NdeI/SalI sites of pGADT7.cDNA fragments of full-length CPK5, N-terminal truncated CPK5(D221A), or C-terminal truncated CPK5 were inserted into the NdeI/EcoRI sites of pGADT7. For split-luciferase assays, a CPK5 cDNA fragment was inserted into the BamHI/SalI sites of pCAMBIA cLuc. cDNA fragments of ABI1, AP2C1, or PP2C6-6 were inserted into the BamHI/SalI sites of pCAMBIA nLuc. For CRISPR/Cas9, a gRNA sequence of STP4 was inserted into the AarI sites of pKIR1.1^40^. All cloning reactions except pKIR1.1 were performed using SLiCE^41^ as described previously. For pKIR1.1, a gRNA fragment was ligated using Ligation high Ver. 2 (TOYOBO). Primers used for plasmid construction are listed in Expanded View Table 1.

### Pathogen inoculation

Alternaria brassicicola (NBRC 31226) was cultured on potato-dextrose agar (PDA) medium and incubated at 25 °C. Single 10 μl drops of spore suspension (5 × 10^5^ spores/ml) were applied to detached 4-week-old Arabidopsis leaves. Inoculated leaves were then incubated on wet paper in petri dishes for 5 days. Six leaf discs from six independent inoculated leaves were pooled into a tube for genomic DNA extraction. The biomass of A. brassicicola on inoculated leaves was assessed by measuring DNA amounts of the fungus (ABU03393) and Arabidopsis (At5g26751)^42^. Strains of Pseudomonas syringae pv. tomato (Pst) DC3000 and Pst DC3000(ΔavrPtoΔavrPtoB) were cultured in NYGA medium (5 g/l Bacto Peptone, 3 g/l yeast extract, 20 ml/l glycerol) with 50 μg/ml rifampicin at 30 °C overnight. The cells were washed twice and resuspended in 10 mM MgCl^2^. These bacteria (OD600 = 0.02) in 0.02% Silwet L-77 were sprayed onto 4-week-old plants, which were then placed under plastic covers for 2 (Pst DC3000) or 3 days (Pst DC3000(ΔavrPtoΔavrPtoB)). Four leaf discs obtained from four independent leaves were pooled. Serially diluted suspensions in 10 mM MgCl_2_ were plated on NYGA agar medium and incubated at 30 °C for 2 days to enumerate colonies.

### In vitro kinase assay

The in vitro kinase assay followed a previously described method^20^ with some modifications. Recombinant proteins were purified from Escherichia coli BL21 (DE3) with TALON Metal Affinity Resin (Takara), Pierce Glutathione Agarose (Thermo scientific), or Amylose Resin (NEB). Kinases (300 ng), their substrates (2000 ng), and phosphatases (1000 ng) were incubated in kinase reaction buffer (100 mM Tris-HCl pH 8.0, 10 mM MgCl_2_, 2 mM EGTA, 1 mM DTT) in the absence or presence of 1.5 mM CaCl_2_ with 100 μM non-radiolabelled ATP containing 370 kBq of [γ-^32^P]ATP (PerkinElmer; NEG002A) for 30 min at 25 °C. Its Ca2+ concentration was estimated at approximately 120 μM by MaxChelator (Ca/Mg/ATP/EGTA Calculator 2.2b) (https://somapp.ucdmc.ucdavis.edu/pharmacology/bers/maxchelator/) To investigate if clade A PP2Cs dephosphorylate kinase substrates, kinase reactions were stopped by addition of 100 μM staurosporine (Wako) for 15 min. Then, clade A PP2Cs were added to the reactions, and incubated for a further 15 min. The reactions were finally stopped by the addition of SDS sample buffer, and incubated at 95 °C for 5 min. The proteins were separated in 10 % polyacrylamide gels. The gels were dried after visualization with Coomassie Brilliant Blue (CBB). Radioactivity in the dried gel was detected by FLA5000 (Fujifilm).

### qRT-PCR analysis

Arabidopsis seedlings were grown on agar medium containing 25 mM sucrose for 6 days, and transferred into 2 ml of liquid medium (Suc 25 mM) in 24-well plates to acclimate plants to liquid medium. The next day, after the medium was replaced with fresh liquid medium with 25 mM sucrose for flg22/chitin treatment or without sugars for 2DG treatment. 1 μM flg22 peptide (QRLSTGSRINSAKDDAAGLQIA, synthesized by Genscript), chitin (250 μg/ml), or 0.5 mM 2DG were applied. Chitin solution was prepared by repeatedly grinding crab or shrimp shell powder (Sigma; C9752) in water with a pestle, and heating to 60 °C. Undissolved particles were removed by centrifugation and filtration. Or, seedlings were grown on agar medium containing 25 mM sucrose for 6 days, and transferred into 2 ml of liquid medium without sugars in 24-well plates to reset cellular sugar conditions. The next day, after changing to fresh medium with the indicated sugars, plants were further incubated for 24 h. Harvested plant samples were frozen and ground in liquid N_2_. Total RNA was isolated using Sepasol RNA I Super G reagent (Nacalai Tesque), followed by treatment with RQ1 RNase-Free DNase (Promega), and reverse-transcribed using ReverTra Ace qPCR RT Master Mix with gDNA remover (TOYOBO) following the manufacturer’s instructions. Quantitative PCR was performed with GoTaq qPCR Master Mix (Promega) on a LightCycler 96 (Roche). The expression levels of genes of interest were normalized relative to those of a reference gene, UBQ10. Relative expression (log_2_) was calculated by subtracting Ct values of genes of interest from those of UBQ10.

### Metabolite quantification

For camalexin and salicylic acid (SA) measurement, frozen and ground samples were homogenized with 500 μl of 80% methanol with 3-indolepropionic acid (TCI; I0032) and d4-SA (Toronto Research; S088127) (0.5 μg each) as internal controls for camalexin and SA, respectively. Mixtures were incubated for 30 min at 65 °C. After centrifugation, 400 μl of the supernatant was transferred to a new tube. Metabolites were repeatedly extracted from the pellets with 80% methanol without internal controls. For sugar measurement, frozen and ground samples were homogenized with 1 ml of extraction buffer (methanol:water:chloroform = 5:2:2) with 10 μg ribitol (Wako) as an internal control. Mixtures were incubated for 30 min at 37 °C. After centrifugation, 900 μl of the supernatant was transferred to a new tube, and 400 μl of water was added. After centrifugation, the upper phase was retrieved. These samples were evaporated using a spin dryer at 50 °C, and subsequently freeze-dried. Samples were sonicated in 30 μl of methoxyamine (Sigma) (20 mg/ml dissolved in pyridine (Wako)), and incubated for 90 min at 30 °C. Subsequently, 30 μl of MSTFA + 1% TMCS (Thermo Fisher Scientific) was added and incubation was continued for 30 min at 37 °C. After centrifugation, the supernatants were subjected to gas chromatography–mass spectrometry analysis. Each sample (1 μl) was separated on a gas chromatograph (7820A; Agilent Technologies) combined with a mass spectrometric detector (5977B; Agilent Technologies). For quantitative determination of metabolites, peaks that originated from selected ion chromatograms (camalexin 272, SA 267, 3-indolepropionic acid 333, d4-SA 271, Glc 319, G6P 387, 2DG 319, 2DG6P 387, ribitol 319) were used.

### Detection of CPK5 phosphorylation

Arabidopsis seedlings were grown on agar medium containing 25 mM sucrose for 6 days, and then transferred into 2 ml of liquid medium with 25 mM sucrose in 24-well plate. The next day, the medium was replaced with fresh liquid medium with 25 mM sucrose for flg22 treatment or without sugars for 2DG treatment. Plants were treated with 1 μM flg22 peptide or 0.5 mM 2DG. In addition, for Glc treatment, seedlings were grown on agar medium containing 25 mM sucrose for 6 days, and transferred into 2 ml of liquid medium without sugars in 24-well plates to reset cellular sugar conditions. The next day, the medium was replaced with fresh medium containing the indicated concentrations of Glc. Plants were incubated for 24 h, then treated with 1 μM flg22. After freezing samples and grinding them in liquid N_2_, proteins were extracted with lysis buffer (50 mM Tris-HCl pH 7.5, 300 mM NaCl, 15% glycerol, 1% Triton X-100, 2 mM DTT). Lambda protein phosphatase (BioAcademia) treatment was for 30 min at 30 °C. Proteins were separated in 8% polyacrylamide gels containing 50 μM Phos-tag Acrylamide (Wako) and 100 μM MnCl2. CPK5-GFP was detected with anti-GFP antibody (B-2, Santa Cruz Biotechnology). Band intensity was measured using ImageJ. For detecting in vitro CPK5 phosphorylation status, HisT7-CPK5 (250 ng) and GST-ABI1 (500 ng) were incubated with reaction buffer (100 mM Tris-HCl pH 8.0, 10 mM MgCl_2_, 2 mM EGTA, 1 mM DTT, 1 mM ATP, 1 mM MnCl2, protease inhibitor cocktail (EDTA-free)(Nacalai)) in the absence or presence of the indicated sugars for 45 min at 30 °C. The reactions were stopped by addition of SDS sample buffer, and incubated at 95 °C for 5 min. The proteins were separated in 10% polyacrylamide gels. CPK5 phosphorylation status was detected using anti-phosphothreonine (p-Thr) antibody (9381S, Cell Signalling Technology). The membranes were stained with CBB.

### Hexokinase activity assay

Hexokinase activity was measured using an enzyme-linked assay following a previously described method^43^ with some modifications. GST-fused recombinant hexokinases were purified from E. coli BL21 (DE3) with Pierce Glutathione Agarose (Thermo Fisher Scientific), and incubated in reaction buffer (50 mM HEPES-NaOH, pH 7.5, 2 mM MgCl_2_, 1 mM EDTA, 9 mM KCl, 1 mM NAD, 1 mM ATP, 2 mM glucose, and 0.8 units NAD-dependent glucose-6-phosphate dehydrogenase (TOYOBO; G6D-321)) at room temperature. Absorbance at 340 nm was monitored continuously with a spectrophotometer (MULTISKAN FC; Thermo Fisher Scientific) to detect NADH production.

### Protein interaction assay

Yeast two-hybrid assays were performed using strain AH109 (Takara). Vectors pGADT7 and pGBKT7 were sequentially transformed into this strain. Transformants were incubated on medium without histidine, leucine, and tryptophan to detect protein interactions. In addition, 1 mM 3-AT was optionally added to medium to investigate strong protein interactions. For split-luciferase assays, Agrobacterium strains (GV3101) transformed with nLuc or cLuc plasmids in infiltration buffer (10 mM MES-KOH pH5.5, 10 mM MgCl_2_) were simultaneously infiltrated into N. benthamiana. leaves. After 2 days, 10 mM luciferin (P1041, Promega) in infiltration buffer was infiltrated into the Agrobacterium-infected leaves. Luminescence was detected using LAS4000 (Cytiva). Luminescence data were coloured using Image J. For pull down assays, MBP-CPK5 (5000 ng) and GST or GST-ABI1 (1000 ng) were added into 100 μl of reaction buffer (100 mM HEPES pH 7.5, 10 mM MgCl_2_, 2 mM EGTA, 2 mM DTT) in the presence/absence of 10 mM of the indicated sugars. After incubation for 30 min at 25 °C, 900 μl of interaction buffer (50 mM Tris-HCl pH 7.5, 150 mM NaCl, 1 mM EGTA, 1mM EDTA, 2 mM DTT, 15 % Glycerol, 1 % Triton-X100) was added to the reactions with MagneGST Glutathione Particles (Promega) in the presence/absence of 10 mM of the indicated sugars. After mixing the tubes at 4 °C for 1 h 15 min, the beads were washed with 1 ml interaction buffer 3 times. SDS sample buffers were added and the tubes were incubated at 95 °C for 5 min. The proteins were separated in 10 % polyacrylamide gels. MBP-tagged and GST-tagged proteins were detected using anti-MBP (E8032S, NEB) and anti-GST antibody (60-021, BioAcademia), respectively.

### Aequorin luminescence measurement

Six-day-old Arabidopsis seedlings expressing p35S-apoaequorin (pMAQ2)^39^ grown on MS agar medium with 25 mM sucrose were transferred into 200 μl of liquid medium with 20 μM coelenterazin h (Wako) in 96-well plates. The next day, after transfer to fresh medium with 20 μM coelenterazin h, luminescence was recorded using a luminometer (LUMINOSKAN; Thermo Fisher Scientific).

### RNA-seq and data analysis

Total RNAs were extracted using Sepazol-RNA I Super G (Nacalai Tesque), followed by treatment with RQ1 RNase-Free DNase (Promega). These DNase-treated RNAs (500 ng) were used for library preparation using BrAD-seq to create strand-specific 3′ digital gene expression libraries^44^. The libraries were sequenced on a HiSeq X Ten platform at Macrogen (Tokyo, Japan), producing 150-base paired end reads. Since the quality of reverse reads was poor due to the poly(A) sequence, only forward reads were used for the analysis. The first 8 bases and adaptors were trimmed and quality filtering was performed by fastp (ver. 0.19.7)^45^. The trimmed and quality-filtered reads were mapped to the Arabidopsis genome (TAIR10) using STAR (ver. 2.6.1b)^46^ and transformed to a count per gene per library using featureCounts (ver. 1.6.0)^47^. Statistical analysis of the RNA-seq data was performed in the R environment (version 3.5.3). Since BrAD-seq involves poly(A) enrichment, mitochondrial and chloroplast genes were excluded. Genes with mean read counts of fewer than 10 per library were considered to be expressed at low levels and were excluded from the analysis. The resulting count data were subjected to trimmed-mean of M-values normalization using the function calcNormFactors in the package edgeR, followed by log-transformation by the function voom in the package limma to yield log_2_ counts per million. To each gene, a linear model was fit by the function lmFit in the limma package with the following terms: Sg = Gg + Rr + ɛg, where S is log_2_ expression value, G is genotype, R is biological replicate and ɛ is residual; or Sgt = GTgt + Rr + ɛgt, where S is log_2_ expression value, GT is genotype:treatment interaction, R is biological replicate and ɛ is residual. For variance shrinkage in the calculation of p-values, the eBayes function in the limma package was used; the resulting p-values were then corrected for multiple hypothesis testing by calculating Storey’s q-values using the function qvalue in the package qvalue. To extract genes with significant expression changes, cutoffs of q-value < 0.01 and |log_2_FC| > 1 were applied. To create heatmaps, average linkage hierarchical clustering with uncentred Pearson correlation as a distance measure was carried out using CLUSTER^48^, followed by visualization using TREEVIEW^48^.

### Data availability

The RNA-seq data used in this study were deposited in the National Center for Biotechnology Information Gene Expression Omnibus database (accession no. GSE169473). Details of the parameters used for quality filtering, mapping and counting are available in the GEO submission.

### Statistics

All data displayed in this study were analysed using R. No statistical methods were used to predetermine sample sizes, but our sample sizes are similar to those generally used in the field. For plant studies, plants were rotated several times in growth chambers to minimize allocation effects. Samples were not blinded for allocation or data analyses. In box plots, centre lines represent the medians, box edges delimit lower and upper quartiles and whiskers show the highest and lowest data points. Experiments in this study were repeated at least three times with similar trends.

## Acknowledgements

We thank Dr. Ryo Matsumoto (Tokushima University) for technical assistance, and Advanced Radiation Research, Education, and Management Centre (Tokushima University) for radioisotope experiments. This work was supported by JST PRESTO (JPMJPR17Q9, K.Y.; JPMJPR17Q6, A.M.), JSPS KAKENHI (20K21280, 21H02157, K.Y.), the Sumitomo Foundation (K.Y.) and the Mitsubishi Foundation (K.Y.).

## Author contributions

**Kohji Yamada**: Conceptualization, Resources, Formal analysis, Data curation, Validation, Funding acquisition, Investigation, Writing-original draft, Project administration. **Akira Mine**: Formal analysis, Data curation, Funding acquisition, Investigation.

## Disclosure and competing interests statement

The authors declare that they have no conflict of interest.

---

**Figure S1.**
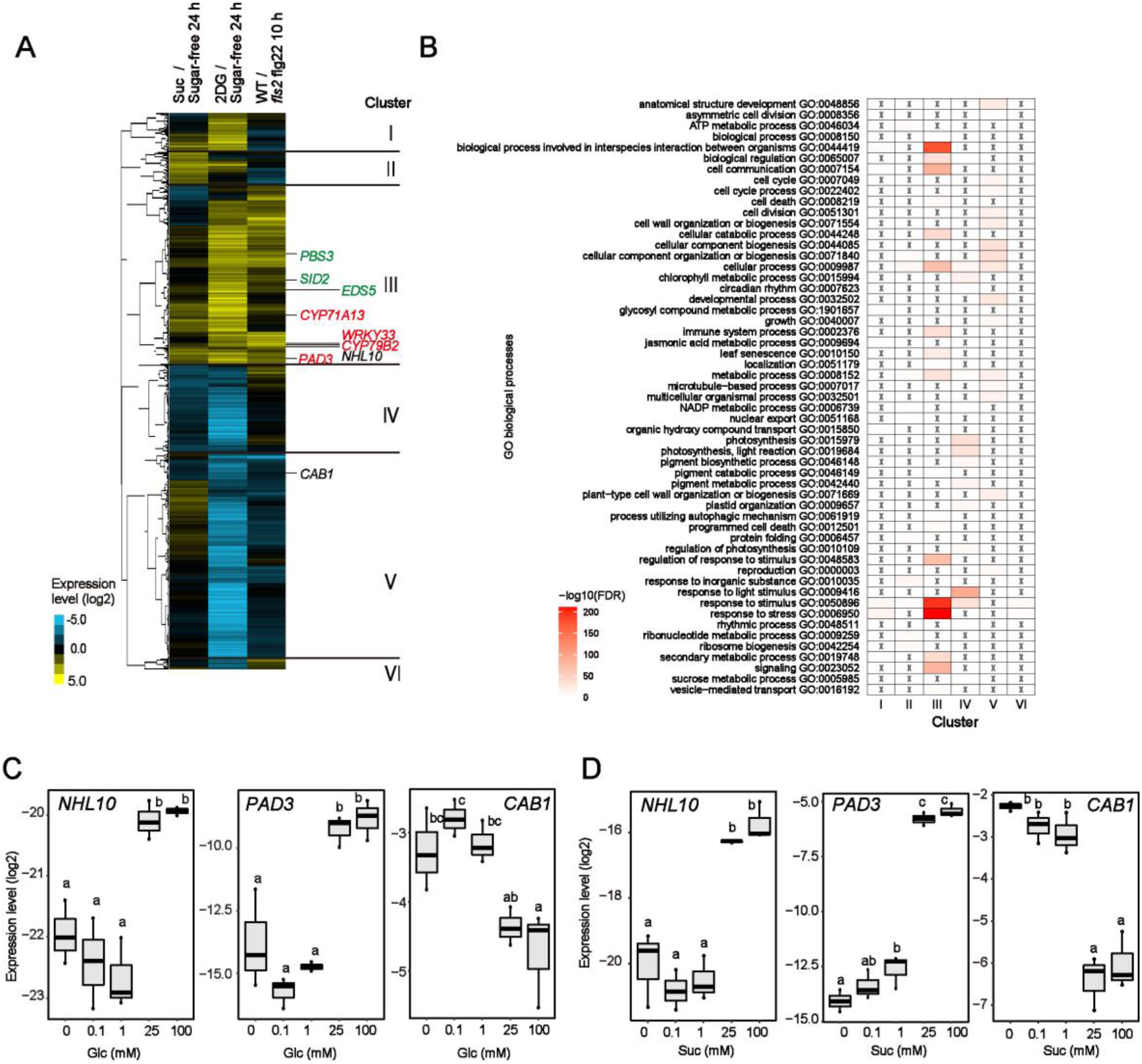
Sugar treatments activate the expression of defence-related genes. **A**, Heat maps displaying expression patterns of genes that show significant expression changes under 1 mM 2DG or 25 mM Suc treatment, relative to sugar-free conditions (q < 0.01 and |log_2_FC| > 1). The log_2_ fold changes were subjected to hierarchical clustering analysis. Camalexin-related genes and SA-related genes were coloured red and green, respectively. **B**, GO analysis of genes in each cluster (C). *P* values were coloured. **C** and **D**, qRT-PCR analysis of the expression of *NHL10* and *CAB1* in *Arabidopsis* seedlings in the presence of Glc (C) or Suc (D) for 24 h (n = 3, biologically independent samples). Individual data points are plotted. Statistically significant differences among samples were determined using one-way ANOVA with multiple comparison tests (Tukey HSD)) and are represented by different letters (*P* <0.05).

**Figure S2.**
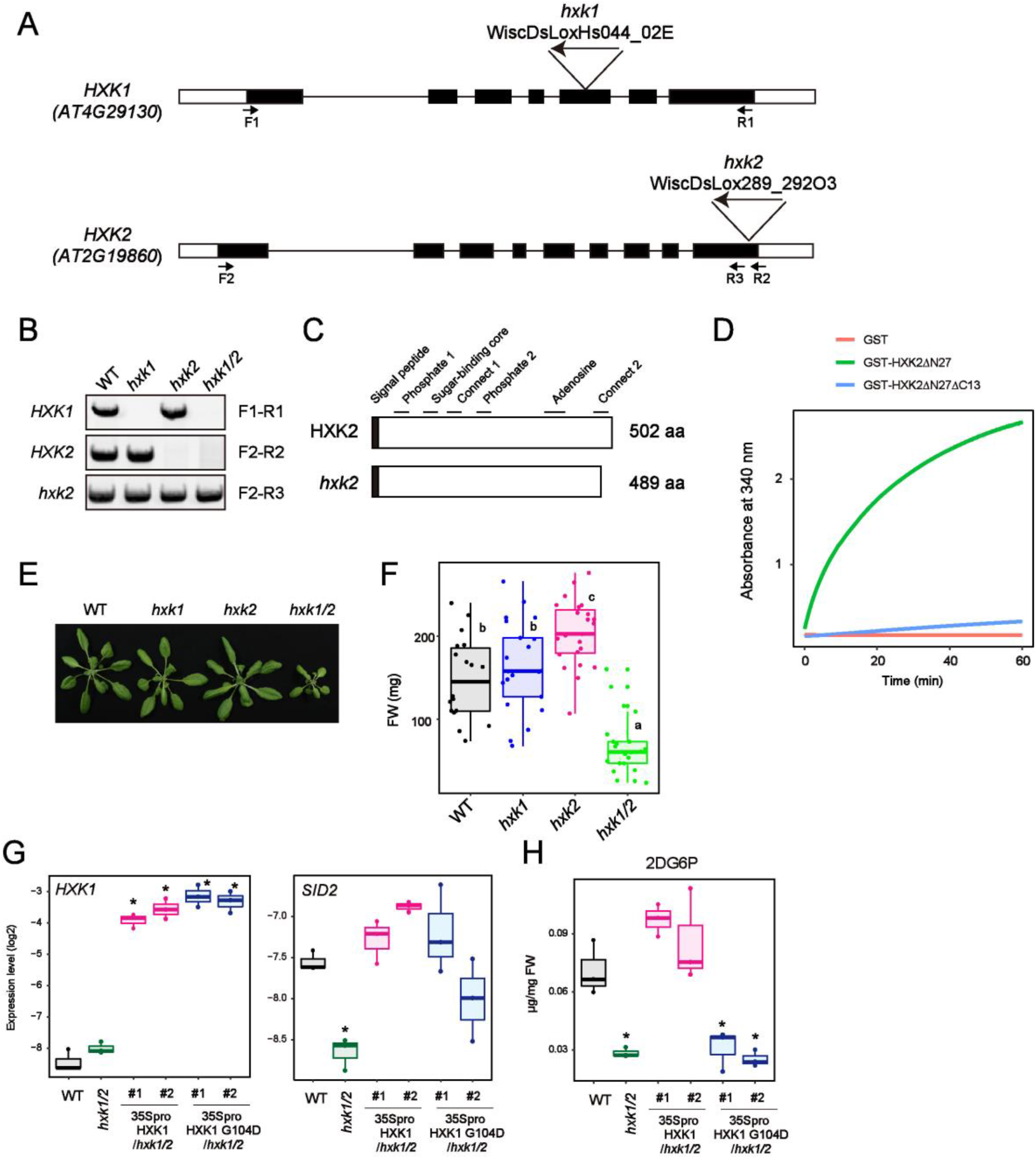
Establishment of *hxk1 hxk2* mutants. **A**, T-DNA insertion mutants of *HXK1* and *HXK2*. Rectangles and lines indicate exons and introns, respectively. Coding regions are coloured black. T-DNA is inserted at each labelled position. **B**, RT-PCR analysis of *hxk1*, *hxk2* and *hxk1/2* mutants. Primers are indicated on the right and their positions are marked in (a). **C**, Presumptive C-terminally truncated HXK2 protein expressed in *hxk2* plants. Conserved regions are labelled above. **D**, Glc phosphorylation activity of HXK2 and 13-aa-deleted HXK2 (mean ± SE, n = 3). Experiments were repeated at least three times with similar trends. **E** and **F**, Growth of *hxk1/2* mutants. **G**, qRT-PCR analysis of defence-related gene expression in *Arabidopsis* seedlings exposed to 0.5 mM 2DG for 8h (n = 3, biologically independent samples). **H**, Quantification of 2DG6P in *Arabidopsis* seedlings 2 h after 0.5 mM 2DG treatment (n = 3, biologically independent samples). FW indicates fresh weight. Statistically significant differences among samples were determined using one-way ANOVA with multiple comparison tests (Tukey HSD (F) or Dunnett’s test between WT and other genotypes (G and H)) and are represented by asterisks or different letters (*P* <0.05).

**Figure S3.**
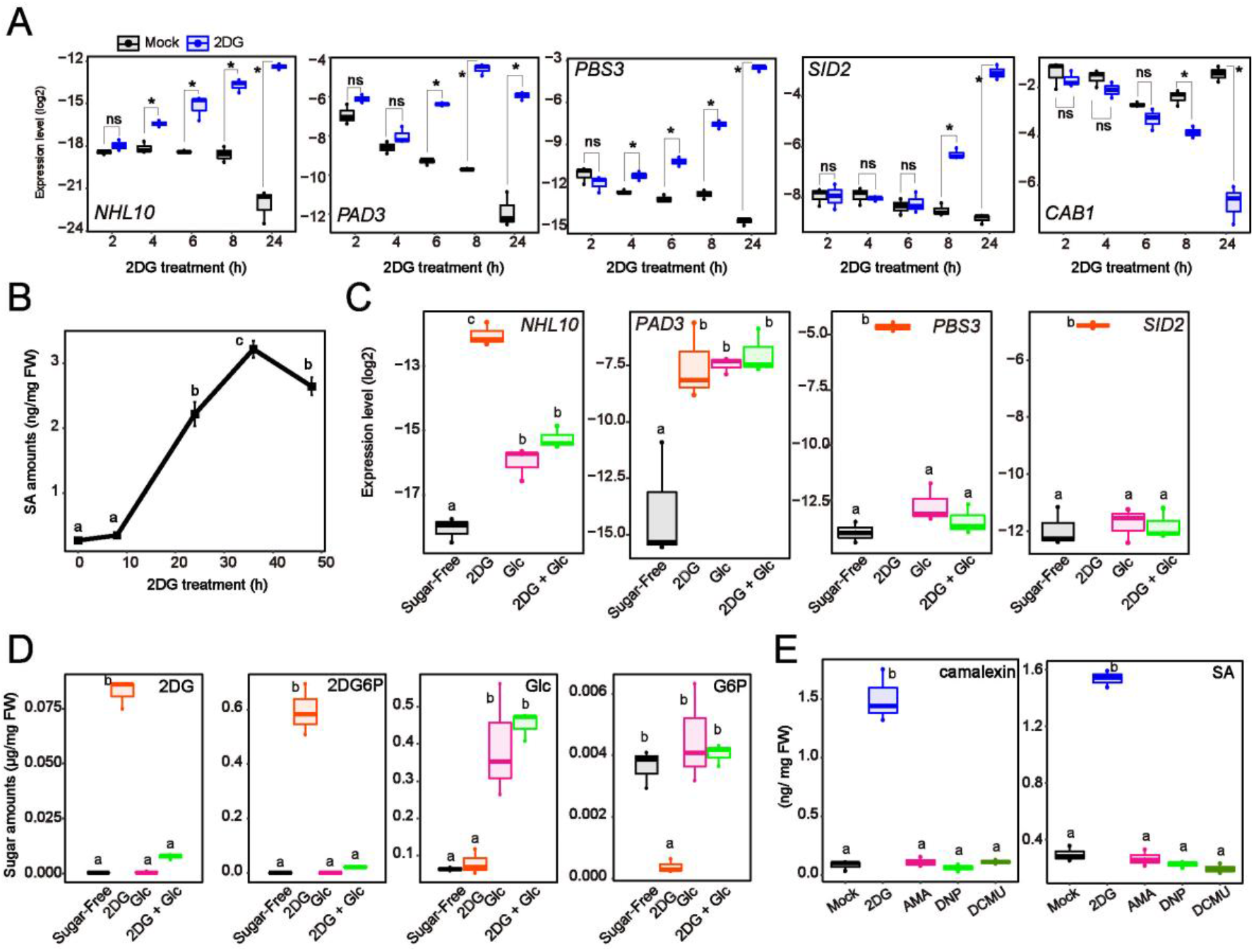
2DG treatment induces defence signalling. **A**, qRT-PCR analysis of defence-related gene expression in *Arabidopsis* seedlings in the presence or absence of 0.5 mM 2DG (n = 3, biologically independent samples). **B**, SA quantification at the indicated time points after 0.5 mM 2DG treatment (mean ± SE, n = 3, biologically independent samples). FW indicates fresh weight. **C** and **D**, qRT-PCR analysis of the expression of defence-related genes (c) and quantification of metabolites (D) in *Arabidopsis* seedlings exposed to 1 mM 2DG and/or 25 mM Glc for 24 h (n = 3, biologically independent samples). 2DG + Glc indicates simultaneous application of 1 mM 2DG and 25 mM Glc. FW indicates fresh weight. **E**, Metabolite quantification 24 h after treatment with 1 mM 2DG, 25 µM AMA, 100 µM DPN, or 40 µM DCMU (n = 3, biologically independent samples). FW indicates fresh weight. Statistically significant differences among samples were determined using the two-tailed Welch’s t test (*P* <0.05) (A) or one-way ANOVA with multiple comparison tests (Tukey HSD) and are represented by asterisks or different letters (*P* <0.05). ns indicates non-significant.

**Figure S4.**
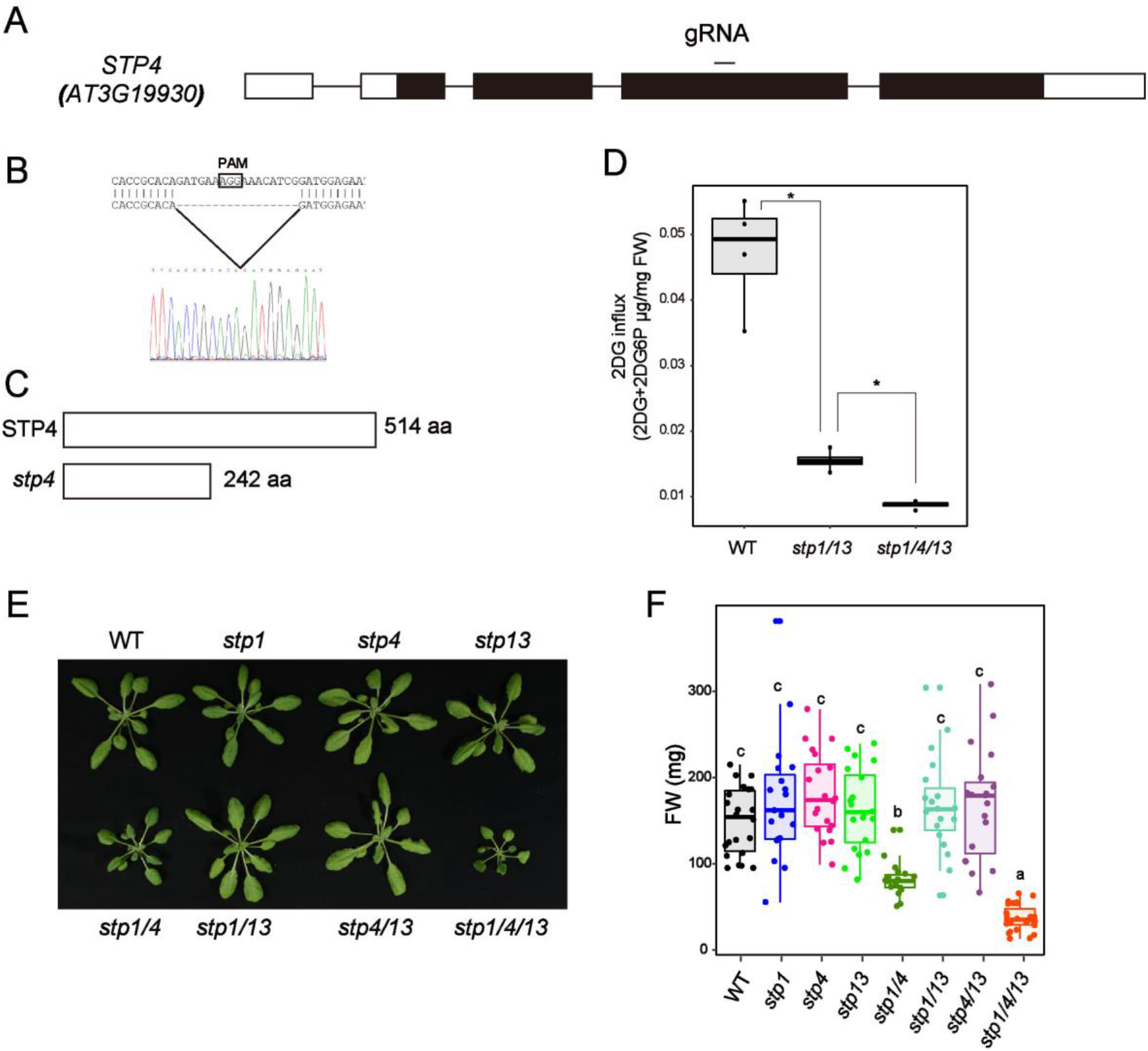
Establishment of *stp1 stp4 stp13* mutants by CRISPR/Cas9. **A**, Genomic structure of *STP4*. Rectangles and lines indicate exons and introns, respectively; the coding region is coloured black. The gRNA for CRISPR/Cas9 was designed at the labelled position. **B**, A 17 bp-deletion occurs in *stp4* mutants. **C**, Predicted length of STP4 proteins in WT and *stp4* mutants. **D**, Glucose influx activity in *Arabidopsis* seedlings 8 h after 1 µM flg22 treatment (n = 3, biologically independent samples). Glucose influx activity was measured by calculating the mixed amounts of 2DG and 2DG6P 1h after 0.5 mM 2DG application. **E** and **F**, Growth of *stp1 stp4 stp13* mutants. Statistically significant differences among samples were determined using the two-tailed Welch’s t test (*P* <0.05) (D) or one-way ANOVA with multiple comparison tests (Tukey HSD) (F) and are represented by asterisks or different letters (*P* <0.05).

**Figure S5.**
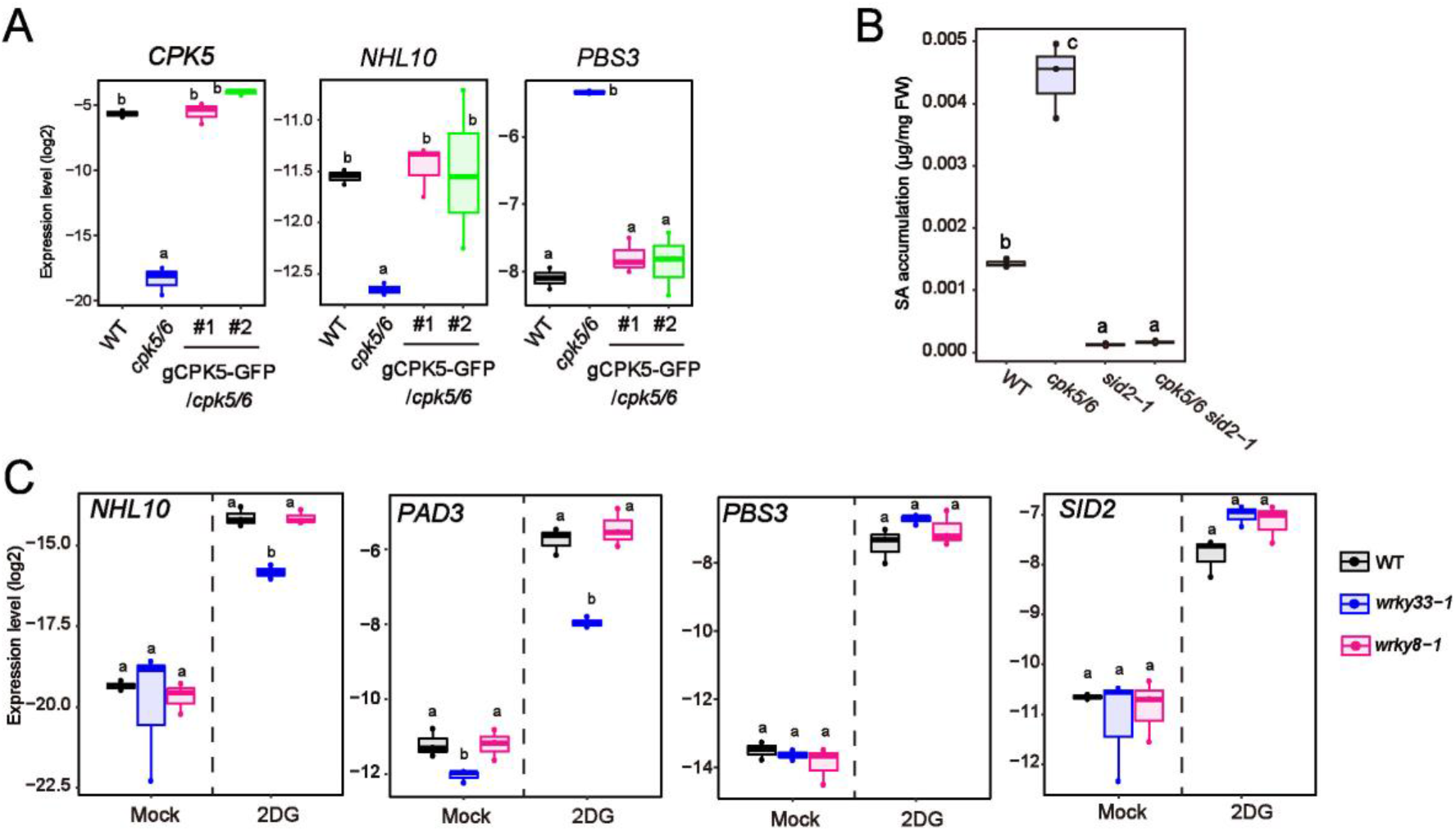
CPK5/6 positively and negatively regulate camalexin and SA synthesis, respectively. **A**, qRT-PCR analysis of defence-related gene expression in *Arabidopsis* seedlings exposed to 0.5 mM 2DG for 8 h (n = 3, biologically independent samples). **B**, SA quantification in *Arabidopsis* seedlings exposed to 0.5 mM 2DG for 24 h (n = 3, biologically independent samples). FW indicates fresh weight. **C**, qRT-PCR analysis of defence-related gene expression in *Arabidopsis* seedlings exposed to mock or 0.5 mM 2DG for 8 h (n = 3, biologically independent samples). Statistically significant differences among samples were determined using one-way ANOVA with multiple comparison tests (Tukey HSD) and are represented by asterisks or different letters (*P* <0.05).

**Figure S6.**
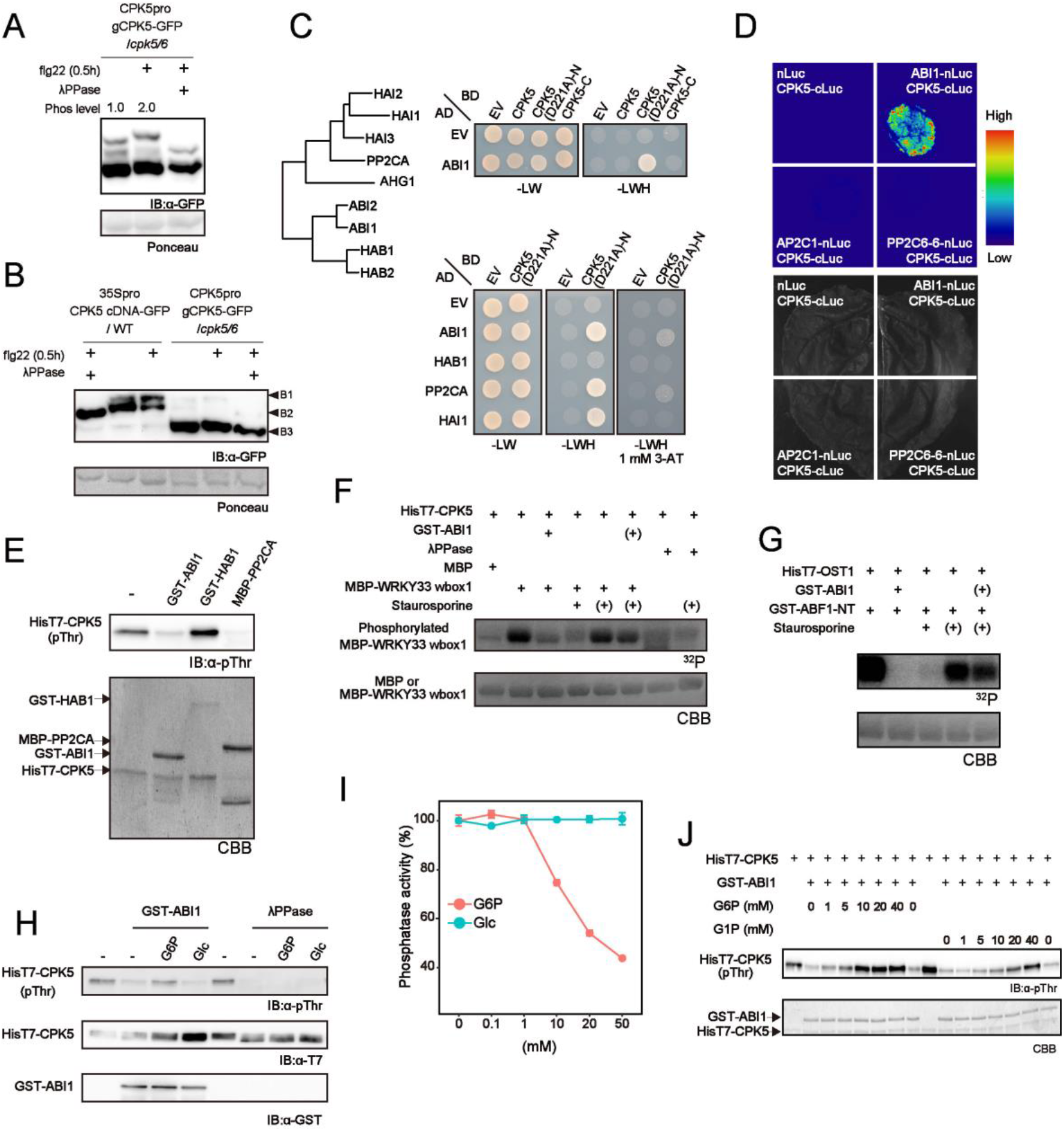
Glucose-6-phosphate suppresses the activity of clade A protein phosphatase 2Cs, leading to elevation of CPK5 phosphorylation. **A**, Immunoblot analysis of CPK5-GFP in *Arabidopsis* seedlings 30 min after 1 μM flg22 application (Mn^2+^-Phos-tag SDS-PAGE). IB denotes immunoblotting with anti-GFP antibodies. Ponceau S-stained membranes are shown below. **B**, Immunoblot analysis of CPK5-GFP in *Arabidopsis* seedlings expressing CPK5(cDNA)-GFP driven by the 35S promoter and a genomic fragment of CPK5 fused with GFP driven by its own promoter 30 min after 1 μM flg22 application (Mn^2+^-Phos-tag SDS-PAGE). IB denotes immunoblotting with anti-GFP antibodies. Ponceau S-stained membranes are shown below. **C**, Protein interactions between CPK5 and clade A PP2Cs by yeast 2-hydrid analysis. The kinase and EF-hand domains are in the N-terminus and C-terminus of CPK5, respectively. CPK5(D221A) is a kinase-dead mutant. Left panel shows a phylogenetic tree for *Arabidopsis* clade A PP2Cs. **D**, Split-luciferase assay for protein interactions between PP2Cs and CPK5 in *N. benthamiana* leaves. **E**, ABI1 and PP2CA, but not HAB1, dephosphorylate CPK5 *in vitro*. IB denotes immunoblotting with anti-pThr antibodies. CBB-stained membranes are shown below. **F** and **G**, Autoradiograph of a WRKY33 wbox1 phosphorylation by CPK5 *in vitro* or an N-terminal ABF1 by OST1 for 30 min with [γ-^32^P] ATP (upper images). Parentheses indicate addition after 15 min. CBB-stained gels are shown below. **H**, λPPase-mediated CPK5 dephosphorylation is not suppressed by G6P. Phosphorylation of CPK5 was detected by anti-phopsho-threonine antibody. 10 mM sugars were applied. IB denotes immunoblotting. **I**, *in vitro* ABI1 phosphatase activity in the presence of G6P or Glc. **J,** ABI1-mediated CPK5 dephosphorylation in the presence of G6P or G1P. Statistically significant differences among samples were determined using one-way ANOVA with multiple comparison tests (Tukey HSD) and are represented by different letters (*P* <0.05).

**Figure S7.**
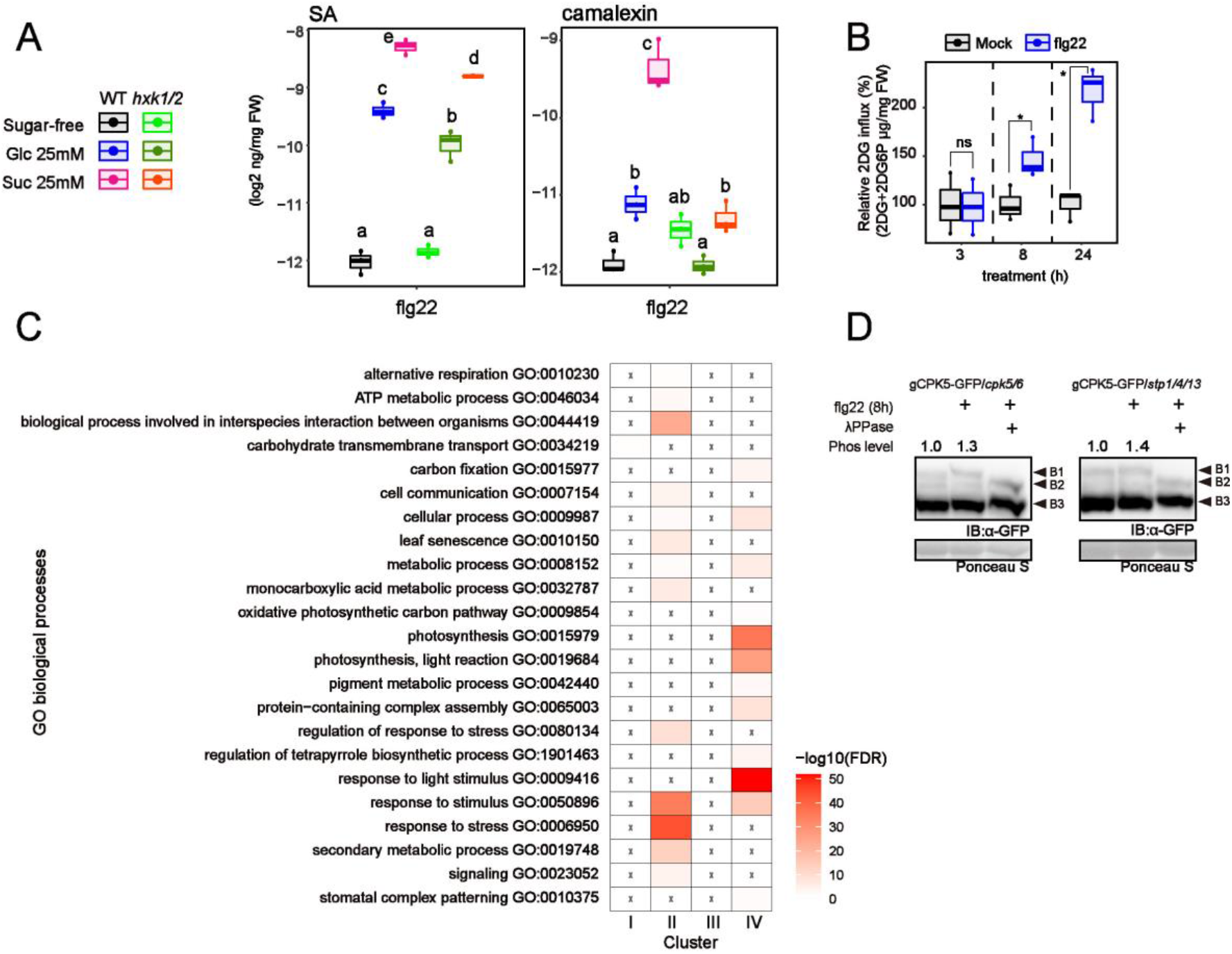
Hexokinase and sugar transporter contribute to flg22-triggered signalling. **A**, Metabolite quantification 8 h after flg22 application in WT and *hxk1/2* plants (n = 3, biologically independent samples). FW indicates fresh weight. **B**, Glucose influx activity in *Arabidopsis* seedlings in response to 1 µM flg22 (n = 3, biologically independent samples). Glucose influx activity was measured by calculating the mixed amounts of 2DG and 2DG6P 1h after 0.5 mM 2DG application. **C**, GO analysis of genes in each cluster (Fig.4c). *P* values were coloured. **D**, Immunoblot analysis of CPK5-GFP in *Arabidopsis* seedlings 8 h after 1 µM flg22 (Mn^2+^-Phos-tag SDS-PAGE). The phospho levels were calculated as ratios of the intensities of B1 and B3. The phospho level of mock 8 h was set to 1.0. IB denotes immunoblotting with anti-GFP antibodies. Ponceau S-stained membranes are shown below. Statistically significant differences among samples were determined using the two-tailed Welch’s t test (*P* <0.05) (B) or one-way ANOVA with multiple comparison tests (Tukey HSD) (A) and are represented by asterisks or different letters (*P* <0.05).

**Figure S8.**
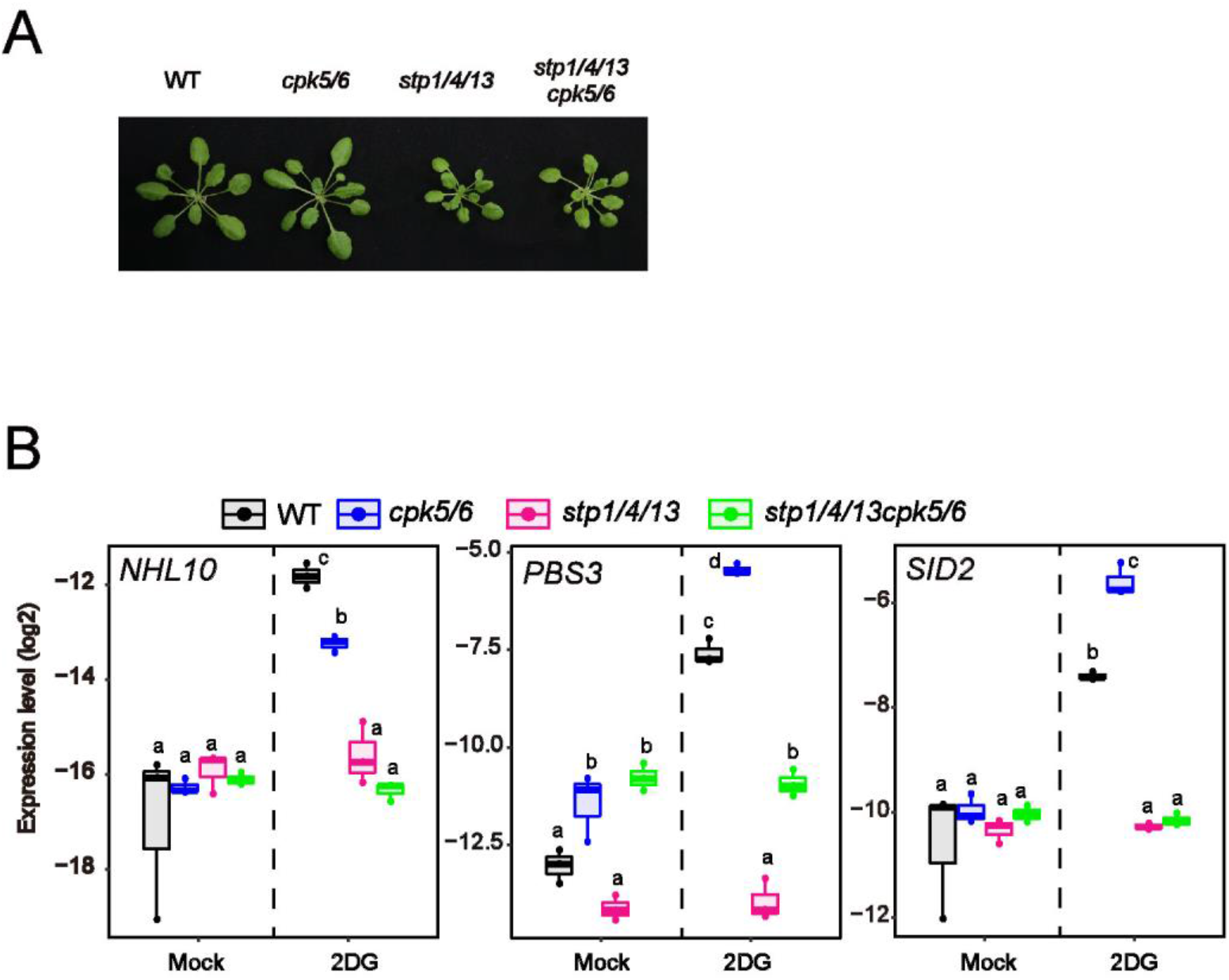
Establishment of *stp1/4/13/cpk5/6* plants. **A**, Growth of *stp1/4/13/cpk5/6* plants **B**, qRT-PCR analysis of defence-related gene expression in *Arabidopsis* seedlings exposed to mock or 0.5 mM 2DG for 8 h (n = 3, biologically independent samples). Statistically significant differences among samples were determined using one-way ANOVA with multiple comparison tests (Tukey HSD) and are represented by asterisks or different letters (*P* <0.05).

**Figure S9.**
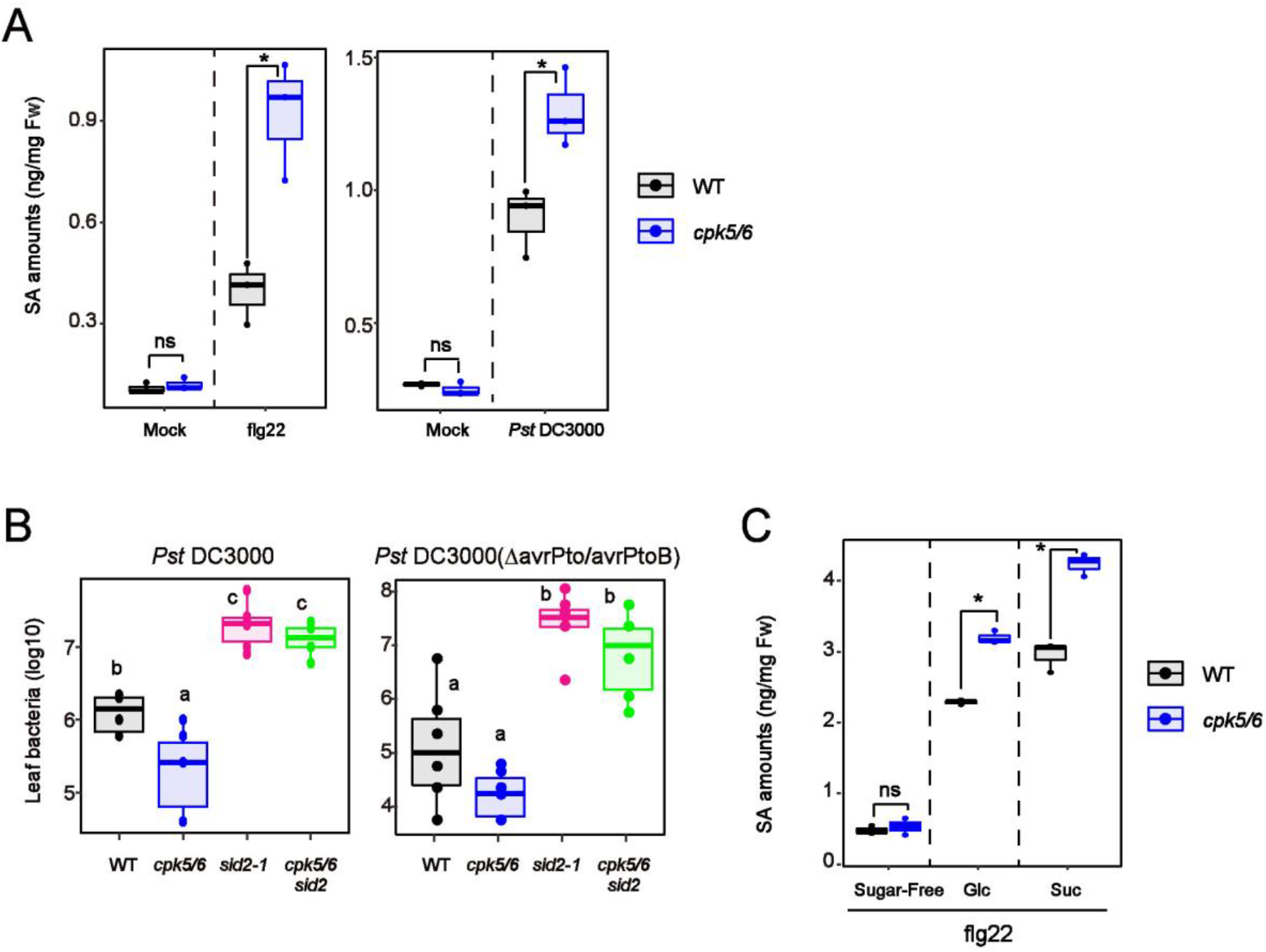
Enhanced SA synthesis decreases susceptibility to pathogenic bacteria in *cpk5/6* plants. **A**, SA quantification in mature leaves of 4-week-old *Arabidopsis* exposed to 1 µM flg22 or syringe-inoculated *Pst* DC3000 for 24 h (n = 3, biologically independent samples). FW indicates fresh weight. **B**, Growth of *Pst* DC3000 (left) or *Pst* DC3000(*ΔavrPtoΔavrPtoB*) (right) spray-inoculated onto rosette leaves of 4-week-old *Arabidopsis*. Bacterial titres were determined 2 (left) or 3 days (right) post-inoculation (n = 6, biologically independent samples). **C**, SA quantification in *Arabidopsis* seedlings exposed to 1 µM flg22 for 8 h in the absence/presence of sugars (n = 3, biologically independent samples). FW indicates fresh weight. Statistically significant differences among samples were determined using the two-tailed Welch’s t test (*P* <0.05) or one-way ANOVA with multiple comparison tests (Tukey HSD) and are represented by asterisks or different letters (*P* <0.05). ns indicates non-significant.

**Figure S10.**
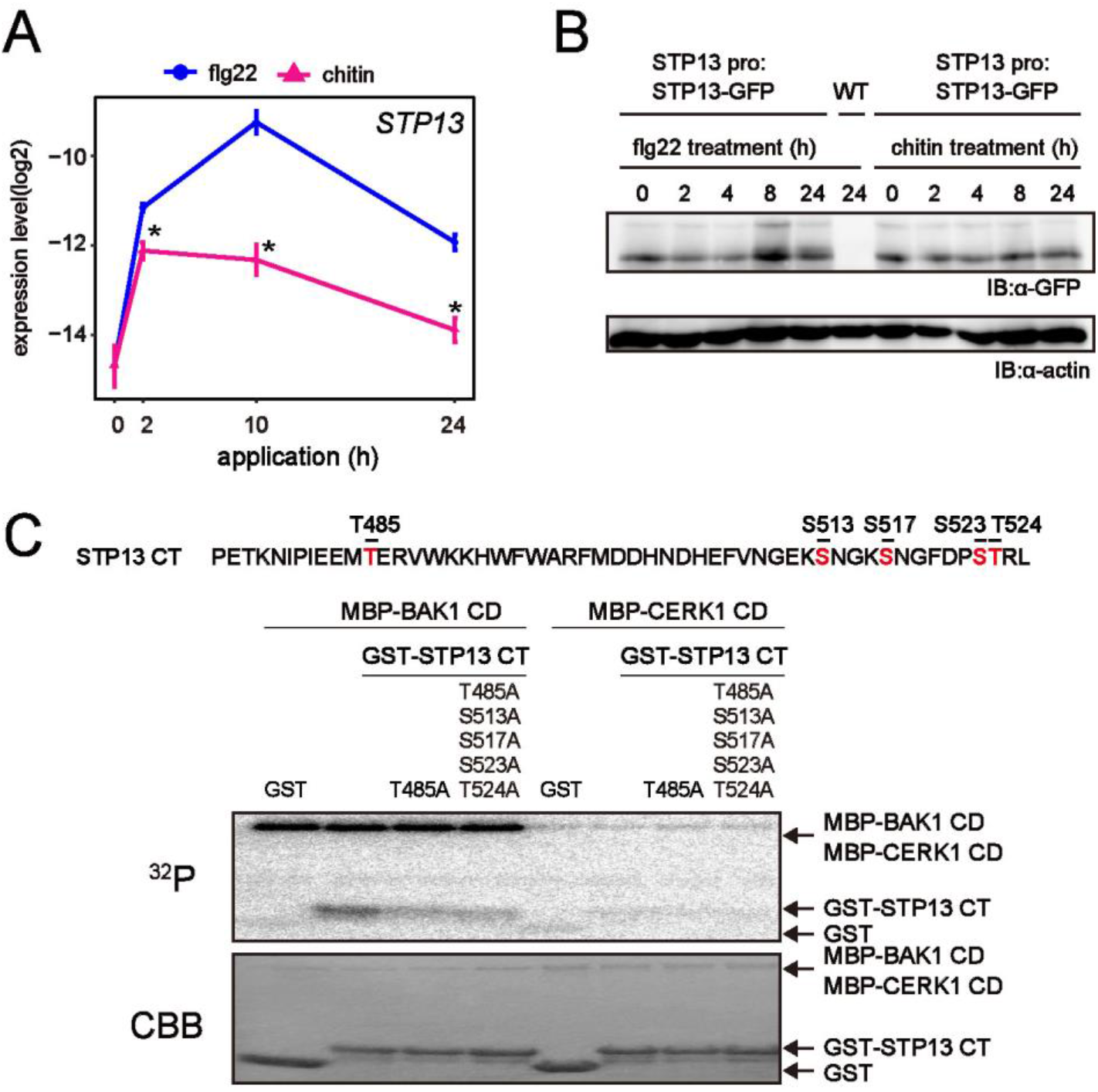
CERK1 does not phosphorylate the T485 residue of STP13. **A**, qRT-PCR analysis of *STP13* expression in *Arabidopsis* seedlings exposed to flg22 or chitin (n = 3, biologically independent samples). **B**, Immunoblot analysis of STP13-GFP in *Arabidopsis* seedlings exposed to flg22 or chitin. IB denotes immunoblotting with anti-GFP or anti-actin antibodies. **C**, Autoradiographs of *in vitro* kinase assays performed by incubating MBP-BAK1 cytoplasmic domain (CD) or MBP-CERK1 CD with GST-STP13 C-terminus (CT) for 30 min with [γ-^32^P]ATP (upper images). CBB-stained gels are shown below. Statistically significant differences among samples were determined using the two-tailed Welch’s t test between flg22 and chitin treatment and are represented by asterisks (*P* <0.05).

## Notes

### Competing Interest Statement

The authors have declared no competing interest.

